# Convergent use of phosphatidic acid for Hepatitis C virus and SARS-CoV-2 replication organelle formation

**DOI:** 10.1101/2021.05.10.443480

**Authors:** Keisuke Tabata, Vibhu Prasad, David Paul, Ji-Young Lee, Minh-Tu Pham, Woan-Ing Twu, Christopher J. Neufeldt, Mirko Cortese, Berati Cerikan, Cong Si Tran, Christian Lüchtenborg, Philip V’kovski, Katrin Hörmann, André C. Müller, Carolin Zitzmann, Uta Haselmann, Jürgen Beneke, Lars Kaderali, Holger Erfle, Volker Thiel, Volker Lohmann, Giulio Superti-Furga, Britta Brügger, Ralf Bartenschlager

## Abstract

Double membrane vesicles (DMVs) are used as replication organelles by phylogenetically and biologically distant pathogenic RNA viruses such as hepatitis C virus (HCV) and severe acute respiratory syndrome coronavirus-2 (SARS-CoV-2). Viral DMVs are morphologically analogous to DMVs formed during autophagy, and although the proteins required for DMV formation are extensively studied, the lipids driving their biogenesis are largely unknown. Here we show that production of the lipid phosphatidic acid (PA) by acylglycerolphosphate acyltransferase (AGPAT) 1 and 2 in the ER is important for DMV biogenesis in viral replication and autophagy. Using DMVs in HCV-replicating cells as model, we found that AGPATs are recruited to and critically contribute to HCV replication and DMV formation. AGPAT1/2 double knockout also impaired SARS-CoV-2 replication and the formation of autophagosome-like structures. By using correlative light and electron microscopy, we observed the relocalization of AGPAT proteins to HCV and SARS-CoV-2 induced DMVs. In addition, an intracellular PA sensor accumulated at viral DMV formation sites, consistent with elevated levels of PA in fractions of purified DMVs analyzed by lipidomics. Apart from AGPATs, PA is generated by alternative pathways via phosphotidylcholine (PC) and diacylglycerol (DAG). Pharmacological inhibition of these synthesis pathways also impaired HCV and SARS-CoV-2 replication as well as formation of autophagosome-like DMVs. These data identify PA as an important lipid used for replication organelle formation by HCV and SARS-CoV-2, two phylogenetically disparate viruses causing very different diseases, i.e. chronic liver disease and COVID-19, respectively. In addition, our data argue that host-targeting therapy aiming at PA synthesis pathways might be suitable to attenuate replication of these viruses.

**One Sentence Summary:** Phosphatidic acid is important for the formation of double membrane vesicles, serving as replication organelles of hepatitis C virus and SARS-CoV-2, and offering a possible host-targeting strategy to treat SARS-CoV-2 infection.

## Main Text

Chronic hepatitis C and COVID-19 are major medical problems. Both diseases are caused by viral infections inflicting a large number of people and having led to millions of deaths ^1, 2^. Chronic hepatitis C is caused by persistent infection with the hepatitis C virus (HCV), while COVID-19 is due to acute infection with the severe acute respiratory syndrome coronavirus-2 (SARS-CoV-2). Both viruses are biologically very distinct e.g. by having a very narrow tropism and a predominantly persistent course of infection in the case of HCV, contrasting the rather broad tropism and acute self-limiting course of infection in the case of SARS-CoV-2. This biological distinction is reflected by their phylogenetic distance with HCV belonging to the *Flaviviridae* and SARS-CoV-2 being a member of the *Coronaviridae* virus family ^3^. In spite of these differences, both viruses possess a single strand RNA genome of positive polarity that is replicated in membranous vesicles in the cytoplasm of infected cells ^4, 5^. These vesicles are induced by viral proteins, in concert with cellular factors, and composed of two membrane bilayers, thus corresponding to double-membrane vesicles (DMVs). These DMVs accumulate in infected cells and can be regarded as viral replication organelle. Viral DMVs have morphological similarity to autophagosomes ^6, 7^, but while autophagy-induced DMVs serve to engulf cellular content and damaged organelles for subsequent degradation, viral DMVs create a conducive and protective environment for productive viral RNA replication. In the case of HCV and SARS-CoV-2, DMVs are derived from the ER ^8, 9, 10^ and can be induced by the nonstructural proteins (NS)3, 4A, 4B, 5A and 5B in the case of HCV ^7^ and the viral proteins nsp3-4 in the case of MERS-CoV and SARS-CoV ^11, 12^, alongside with co-opted host cell proteins and lipids. Here, we set-out to search for common host cell factors exploited by the phylogenetically distant HCV and SARS-CoV-2 to build up their cytoplasmic replication organelle.

Using HCV as a model to study DMV biogenesis, we purified DMVs under native conditions and determined their molecular composition by proteomic profiling (Fig. 1A and B). To this end we used human hepatoma cells (Huh7) containing a self-replicating HCV replicon RNA (designated sg4B^HA^31R; ^13^) in which NS4B was HA-tagged (fig. S1A). This RNA replicates autonomously and induces an extensive array of DMVs that can be isolated by HA-affinity purification ^13^. Mass spectrometry-based proteomics analysis identified a total of 1487 proteins significantly enriched in the NS4B-HA sample relative to the untagged technical negative control (using SAINT average *P*-values >0.95) (data S1). Label free quantitation (LFQ) revealed a major overlap of proteins (1542) between the NS4B-HA complex and HCV-naïve ER membranes purified in parallel from Huh7 cells stably expressing HA-tagged Calnexin (CNX-HA) (Fig. 1B and fig. S1B). Of note, 309 proteins were significantly enriched in the NS4B-HA sample relative to the ER control with an over-representation of proteins involved in RNA metabolism, intracellular vesicle organization and transport as well as endomembrane organization (fig. S2). Given our interest in identifying proteins of relevance for DMV formation, we selected 139 candidates with a bias for proteins involved in vesicle transport and biogenesis as well as lipid metabolism. These candidates were validated with respect to their role in HCV replication by using RNA interference-based screening (Fig. 1C and data S2). In this way we could validate 38 hits as HCV dependency factors. Amongst identified hits were acylglycerolphosphate acyltransferase (AGPAT) 1 and 2, two enzymes that catalyze the *de novo* formation of phosphatidic acid (PA), a precursor to di- and triacylglycerols as well as all glycerophospholipids ^14, 15^. In addition, PA is involved in signaling and protein recruitment to membranes and, owing to its small and highly charged head group, promotes membrane curvature ^16, 17, 18^. Since these properties might be involved in DMV formation, we focused our subsequent analysis on AGPATs.

**Fig. 1.**
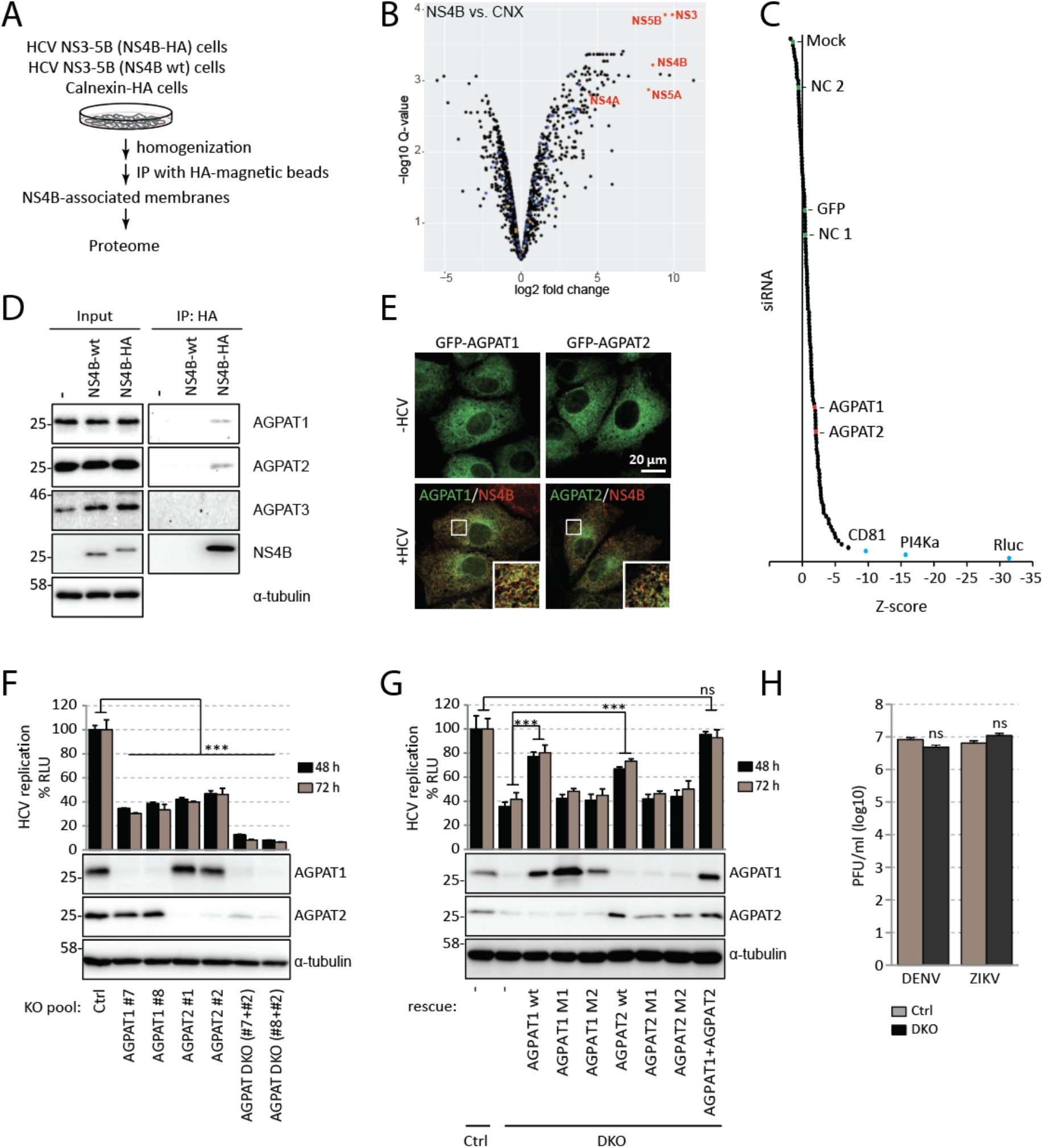
Proteome analysis of HCV-induced DMVs identifies AGPATs as host dependency factors critically contributing to viral replication. (A) Experimental approach used to purify DMVs from HCV-replicating cells. (B) Volcano plot of differentially enriched interactors of NS4B and calnexin (CNX). Q-values were calculated using the limma software package and corrected for multiple hypothesis testing. Viral proteins are highlighted with red letters. A magnified view with protein hits labeled is given in fig. S1B. (C) A total of 139 genes were selected from the DMV proteome and validated by siRNA screening (3 siRNAs per gene). CD81, PI4KA and Rluc were used as positive controls; NC (negative control)1, NC2, GFP and mock infection served as negative controls. A summary of the screening is given in Data S2. (D) Endogenous AGPAT1 and 2, but not AGPAT3 are contained in NS4B-associated membranes. Membranes were purified from naïve Huh7-Lunet cells (-), or Huh7-Lunet cells containing a subgenomic replicon without or with an HA-tag in NS4B (NS4B-wt and NS4B-HA, respectively). Captured proteins were analyzed by western blot, along with the input (2%). α-tubulin served as loading control. (E) Colocalization of NS4B with AGPAT1 and 2. Huh7-Lunet cells stably expressing AGPAT1- or AGPAT2-GFP were transfected with *in vitro* transcripts of the HCV genome Jc1 and fixed 48 h post-transfection. (F) Effect of AGPAT KO on HCV replication. Huh7.5 cells were infected with lentiviruses encoding AGPAT-targeting-sgRNA and 5 days later, infected with an HCV reporter virus (JcR2a). After 48 h and 72 h, *renilla* luciferase activities in cell lysates, reflecting viral RNA replication, were quantified. Graph shows average and SD from 3 independent experiments. Significance was calculated by a paired t-test. ***, p<0.001. Abundance of AGPAT proteins is shown on the bottom. α-tubulin served as loading control. (G) Enzymatic activity of AGPAT is required for HCV replication. KO cells were reconstituted with sgRNA-resistant AGPAT wild-type (wt) or catalytically dead mutants (M1 and M2) by lentiviral transduction. Cells were infected with JcR2a, and *renilla* luciferase activities were quantified. Graph shows average and SD from 3 independent experiments. Significance was calculated by paired t-test. ***, p<0.001. ns, p>0.05. Note the complete rescue by AGPAT1 and 2 co-expression. Abundance of AGPAT proteins is shown below the graph; α-tubulin served as loading control. (H) AGPAT1/2 DKO does not affect DENV or ZIKV propagation. Cells were infected with DENV-2 (strain 16681) or ZIKV (strain H/PF/2013) and 48 h later virus titer was quantified by plaque assay. Graph shows the average and SD from 3 independent experiments. Significance was analyzed by a paired t-test. ns, p>0.05. PFU, plaque forming units.

AGPATs play crucial roles in lipid homeostasis, because enzyme-inactivating mutations in AGPAT2 are linked to congenital generalized lipodystrophy and defects in PA metabolism as well as autophagy are associated with neurological disorders and chronic obstructive pulmonary disease ^18, 19^. Moreover, severe lipodystrophy as well as extreme insulin resistance and hepatic steatosis have been observed in AGPAT2^-/-^ mice ^14^. To date, 11 AGPATs have been identified in mammalian cells. AGPAT1 to 5 preferentially utilize lysophosphatidic acid (LPA) as an acyl donor while AGPAT6 to 11 preferentially utilize alternative lysophospholipid substrates or have a preference for glycerol-3-phosphate. Thus, only AGPAT1 to 5 function as true LPA acyltransferases ^14^. To establish which AGPAT family members are found in NS4B-associated membranes, FLAG-tagged versions of each of the 5 AGPATs were transiently expressed in cells containing the HCV replicon sg4B^HA^31R (fig. S3A). Pull-down of NS4B-HA revealed association with AGPAT1 and 2, and to a lesser extent with AGPAT3, but not with AGPAT4 and 5. Additionally, endogenous AGPAT1 and 2 were detected in NS4B-HA containing membranes isolated from replicon-containing cells (Fig. 1D), whereas AGPAT 3 was not enriched. Moreover, in HCV infected cells AGPAT1 and 2 were recruited to NS4B-containing sites that most likely correspond to sites of DMV accumulation ^13^ (Fig. 1E).

To validate the role of AGPAT1 and 2 in HCV replication, we created knock-out cells using CRISPR/Cas9. Although we observed reduced cell growth of stable double knock-out (DKO) cells 8 days after transduction of guide RNAs, single KO cell pools showed no decrease in cell growth and could be used for transient knock-out of the other AGPAT gene without impacting cell viability for up to 8 days after transduction (fig. S3B). Using this approach, we observed that AGPAT1/2 DKO impaired lipid droplet formation (fig. S3, C to E) as shown previously ^20, 21^, confirming disruption of AGPAT1/2 function. To monitor the impact of single KO and AGPAT1/2 DKO on HCV replication, cells were infected with an HCV reporter virus and viral replication was determined by using luciferase assay. While single KO suppressed HCV replication by ~50-70%, a reduction by ~90% was observed in DKO cells (Fig. 1F). Even stronger replication suppression was observed with a subgenomic replicon (fig. S4A), confirming that AGPAT depletion affected viral RNA replication and not virus entry or assembly. Of note, replication was completely restored by stable expression of AGPAT1 and 2 in DKO cells, which was not the case with either or both enzymatically inactive mutants (Fig. 1G). In contrast, replication of Dengue virus (DENV) and Zika virus (ZIKV), also belonging to the *Flaviviridae* family, but inducing morphologically different membrane alterations, i.e. ER membrane invaginations ^4^, was not affected as determined by plaque assay or with a reporter virus (Fig. 1H and fig. S4B, respectively). These results suggest that enzymatically active AGPAT1 and 2 are required for HCV replication with both AGPATs having partially redundant functions.

Next, we determined the impact of AGPAT KO on HCV-induced DMV formation. Since AGPAT1/2 DKO reduces RNA replication, we employed a replication-independent system in which DMV production is induced by the sole expression of an HCV NS3-5B polyprotein fragment that undergoes self-cleavage to produce functional NS3, 4A, 4B, 5A and 5B ^8, 22^ (Fig. 2A). To determine the replicase subcellular location by fluorescence microscopy, NS5A was fluorescently tagged with EGFP. This tagging has no effect on replicase functionality ^8, 22^. While expression of this polyprotein induced a high number of DMVs in control cells, DMV abundance was dramatically reduced in AGPAT1/2 DKO cells (Fig. 2, A and B), although amounts of viral proteins were comparable in control and DKO cell pools (Fig. 2C). Moreover, DMVs had a smaller diameter in AGPAT2 KO cells (fig. S4C). These results argue for a pivotal role of AGPATs in HCV DMV biogenesis.

**Fig. 2.**
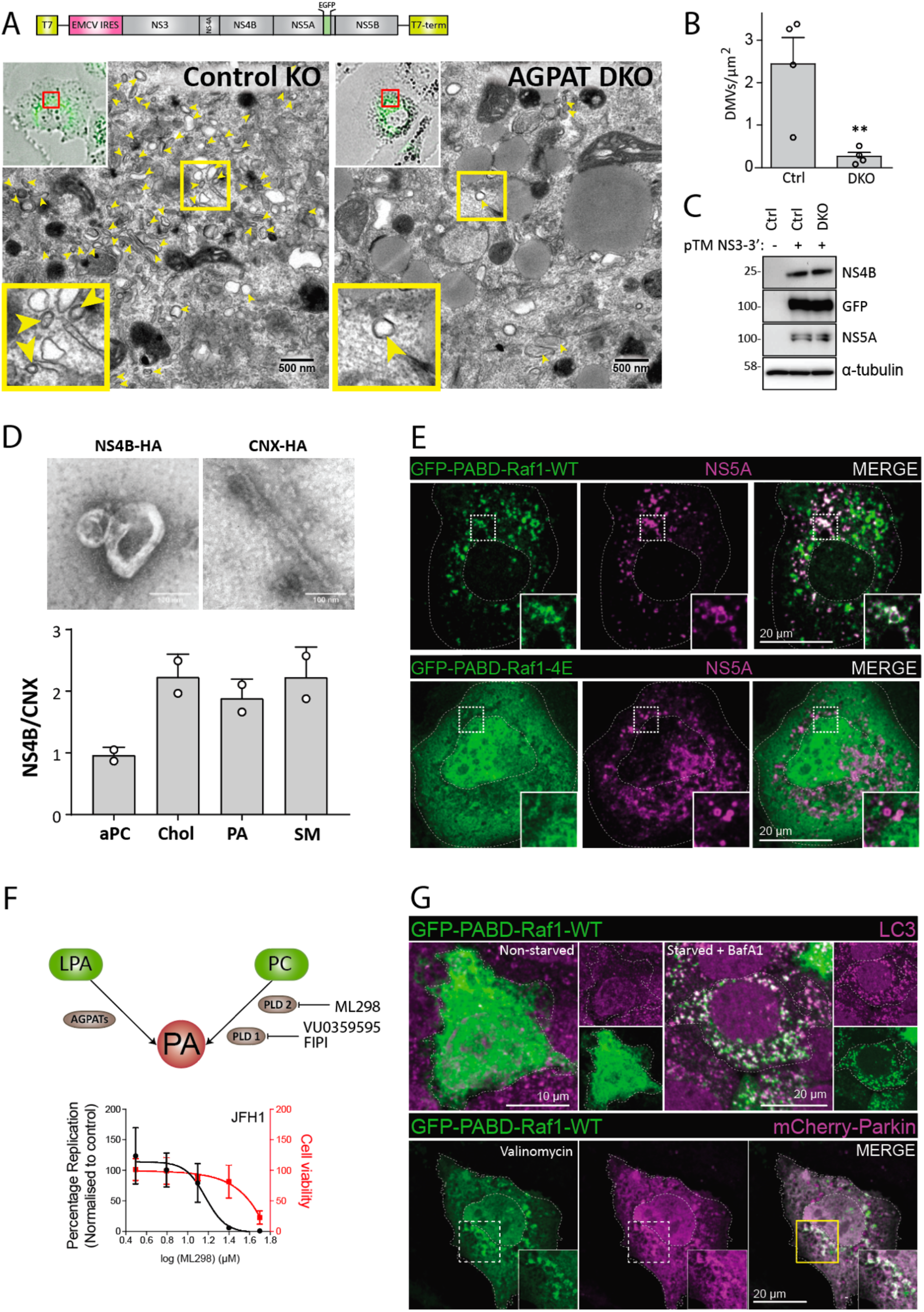
Requirement of AGPATAs for HCV-induced DMV formation and PA accumulation on HCV-induced DMVs and autophagy-related structures. (A to C) AGPAT1/2 DKO dampens DMV formation induced by HCV. Huh7-derived cells stably expressing the T7 RNA polymerase and containing or not a double knock-out (DKO) of AGPAT1 and 2 were transfected with a HCV replicase-encoding plasmid containing a GFP insertion in NS5A (construct pTM NS3-3’/5A-GFP, top panel). Transcripts are generated from the plasmid in the cytoplasm via the T7 promoter and terminator (T7-term) sequence and the HCV NS3 – 5B coding region is translated via the IRES of the encephalomyocarditis virus (EMCV). (A) After 24 h, cells were fixed and subjected to CLEM. Low resolution confocal microscopy images identifying transfected cells are shown on the top left. The area in the red box is shown in the corresponding EM image. Yellow arrow heads indicate DMVs. Insets at the bottom indicate zoomed-in regions. (B) DMVs within whole cell sections were counted and divided by cell area (μm^2^). Graph shows average and SD from 4 different transfected cells. Cells expressing comparable level of HCV replicase were selected for EM analysis. Significance was calculated by a paired t-test. **, p<0.01. (C) Expression levels of NS4B and NS5A in transfected cells were determined by western blotting. (D) Lipidome analysis of HCV-induced DMVs. Extracts of Huh7 cells containing the subgenomic replicon sg4B^HA^31R (NS4B-HA) and Huh7 cells stably overexpressing HA-tagged Calnexin (CNX-HA) and control Huh7 cells were prepared as described in supplementary methods and used for HA-affinity purification under native conditions. An aliquot of the sample was analyzed by electron microscopy (top panels) whereas the majority was subjected to lipidome analysis by using mass spectrometry. Values obtained for the NS4B-HA sample were normalized to those obtained for the CNX-HA sample that was set to one. The complete list of analyzed lipids is summarized in data S3. (E) PA accumulation at NS5A containing structures. Huh7-Lunet/T7 cells were transfected with a construct analogous to the one in panel A, but containing a mCherry insertion in lieu of GFP, along with an EGFP-tagged wildtype (WT) or mutant (4E) PA sensor (construct pTM-EGFP-PABD-Raf1-WT or -4E). Twenty-four hours later, GFP-PABD and NS5A-mCherry were visualized by fluorescence microscopy. White boxes indicate regions magnified in the lower right of each panel. (F) Top panel: Alternate PA biosynthesis pathways via lysophosphatidic acid (LPA) or phosphatidylcholine (PC) catalyzed by AGPATs or PLDs, respectively. Bottom panel: Huh7-Lunet/T7 cells were electroporated with in vitro transcripts of a subgenomic HCV reporter replicon encoding the firefly luciferase. Four hours after transfection, different concentrations of PA synthesis inhibitors were added to the cells and luciferase activities were analyzed at 48 h after electroporation. Graph shows average and SD from 3 independent experiments. Cell viability determined by CellTiter-Glo luminescent assay is indicated with the red line. (G) PA recruitment to autophagy-related structures in selective and non-selective autophagy. Top panel: Huh7-derived cells expressing EGFP-PABD-Raf-1 were incubated in growth medium (top left panels) or in serum-free medium with 200 nM BafA1 (top right panels) for 3 h. Cells were fixed and stained with a LC3 specific antibody. Bottom panel: For selective autophagy, mCherry-tagged Parkin was co-expressed with EGFP-PABD-Raf1, followed by incubation with 10 μM Valinomycin to induce mitophagy. Cells were fixed after 3 h, and GFP-PABD and mCherry-Parkin were visualized by fluorescence microscopy. Images in panels E and G are maximum intensity projections.

Given that AGPAT1 and 2 are important for DMV formation and their enzymatic activity is required for HCV replication, we next focused on their reaction product, i.e. the lipid PA. To quantify the amount of PA associated with HCV-induced DMVs and compare it to ER membranes, we determined the lipidome of highly purified DMVs isolated from cells containing the sg4B^HA^31R replicon (Fig. 2D). Consistent with earlier results, these membranes contained elevated amounts of cholesterol and sphingolipids, which served as positive controls, relative to ER membranes purified in parallel ^13, 23^. Of note, PA abundance in DMVs also was increased in comparison to ER membranes, whereas the level of diacyl phosphatidylcholine (aPC) and several other lipids was not affected (Fig. 2D; for further lipids see data S3).

To confirm these findings in single cells, we used two alternative methods to detect PA by fluorescence microscopy. First, we generated a recombinant protein composed of GST that was fused to the PA binding domain (PABD) derived from yeast Spo20p (fig. S5A and B). As a specificity control we employed the analogous sensor protein containing a mutation in the PABD that abolishes PA binding, and GST alone ^24^. These proteins were introduced via transient permeabilization into Huh7 derived cells (fig. S5C). In cells treated with phorbol 12-myristate 13-acetate (PMA), a potent activator of phospholipase D-mediated PA production, as expected the intact sensor predominantly stained the plasma membrane, which was not the case with the PA non-binding mutant or GST alone, confirming specificity of the signal (fig. S5D). Moreover, also in cells that were not treated with PMA, the PA sensor predominantly stained the plasma membrane (fig. S5D, right panel). Using this assay, we monitored intracellular PA distribution in HCV replicon-containing cells and observed PA colocalization with NS4B (fig. S5E). As second assay for intracellular PA detection, we created a GFP-tagged sensor fused to the PABD of Raf1, a serine-threonine kinase recruited to cellular membranes via its interaction with Ras and PA ^25^. While in control Huh7 cells this PA sensor displayed a diffuse pattern (fig. S6A), upon co-expression of the HCV NS3-5B polyprotein the sensor accumulated in NS5A-positive puncta (Fig. 2E). Of note, a control PA sensor containing mutations in the PABD of Raf1 (mutant 4E) ^26^ displayed only a diffuse pattern in NS3-5B expressing cells (Fig. 2E), supporting specificity of the signal and PA recruitment to HCV replication sites.

Since these data suggest an important role of AGPAT1 and 2-dependent PA enrichment on HCV-induced DMVs, we hypothesized that other pathways contributing to PA generation in cells might also play a role in HCV replication. Apart from AGPATs, one other route for PA synthesis is through hydrolysis of phosphatidylcholine (PC) by phospholipase D1 (PLD1) and D2 (PLD2) enzymes (Fig. 2F, top panel) ^17, 27^. To test the role of PLD1/2 enzymes in HCV replication, we employed a pharmacological approach using 3 different PLD1/2 inhibitors. Treatment with PLD2 inhibitor ML298 caused replication inhibition at a concentration that did not significantly reduce cell viability (~25 μM; Fig. 2F, bottom panel), whereas for the other drugs the reduction in HCV replication correlated with cytotoxicity (not shown). In summary, these results suggest that PA generated via AGPAT1/2, and possibly by alternative PA synthesis pathway, contributes to HCV replication by supporting the formation of DMVs, which is the site of viral RNA amplification.

Virus-induced DMVs are morphologically analogous to autophagosomes generated during autophagy ^7^; therefore, we tested if PA would be recruited to and is required for autophagy-induced DMVs. To this end, we monitored the localization of the GFP tagged PA sensor with markers for DMVs induced during nonselective and selective autophagy. To monitor DMV formation induced during nonselective autophagy, cells were incubated in starvation medium with or without bafilomycin A1 (BafA1), an inhibitor of the vacuolar-type H^+^-ATPase inducing the accumulation of LC3-positive puncta, which are indicative of autophagosomes. For selective autophagy events, we focused on the induction of DMVs during mitophagy induced by treatment of the cells with valinomycin (Val) ^28, 29^. As shown in Fig. 2G (top row), the PA sensor GFP-PABD-Raf1 was rather uniformly distributed throughout the cell in non-induced cells. However, induction of nonselective autophagy by serum starvation led to a significant increase in the number of LC3 puncta with GFP-PABD-Raf1 relocalizing to these puncta (Fig. 2G). Similarly, induction of mitophagy by Val treatment caused an abundant association of mCherry-Parkin puncta with GFP-PABD-Raf1 (Fig. 2G, lower panel), whereas in control cells not treated with Val, no such association was found (fig. S6B). Next, we investigated the functional role of PA generation during nonselective and selective autophagy. Consistent with the relocalization of PA to LC3 puncta during nonselective autophagy, PA inhibitors targeting PLD1, PLD2 and AGPATs, applied as short-term treatments and at non-toxic concentrations, significantly reduced the accumulation of LC3 puncta (fig. S7). These findings are consistent with a recent study suggesting that PA generated on the ATG16L1-positive autophagosome precursor membrane contributes to autophagosome formation ^30^. Of note, a third pathway for PA production via phosphorylation of diacylglycerol (DAG) by diacylglycerol kinase (DAGK) ^27^, did not contribute to PA accumulation or increase in LC3 puncta during nonselective autophagy (fig. S7).

Having found that AGPAT1 and 2, and their reaction product PA, are involved in DMV formation induced upon HCV infection and in, morphologically similar, DMVs generated during autophagy, we hypothesized that AGPATs and PA might also be involved in the biogenesis of replication organelles of other unrelated RNA viruses, e.g., coronaviruses, which also utilize DMVs as viral replication sites ^9, 10^. Hence, we investigated the role of AGPATs in the DMV biogenesis of SARS-CoV-2, the causative agent of the ongoing COVID-19 pandemic. In the first set of experiments, we studied the recruitment of AGPATs to SARS-CoV-2 induced DMVs. In the case of MERS-CoV and SARS-CoV, formation of DMVs with structural resemblance to those observed in infected cells can be induced by the sole expression of viral nonstructural protein (nsp)3-4, which is an ~270 kilodalton large polyprotein fragment undergoing self-cleavage ^12^. Building on these results we first determined whether the same applies to SARS-CoV-2. Huh7-derived cells stably expressing T7 RNA polymerase were transiently transfected with a T7 promoter driven SARS-CoV-2 HA-nsp3-4-V5 expression construct or the empty vector (fig S8A). Using immunofluoresence with an HA-specific antibody in many cells we observed clusters of HA-nsp3 (fig. S8B). Western blotting confirmed efficient self-cleavage between nsp3 and nsp4 (fig. S8C). To identify membrane alterations in HA-nsp3-4-V5 expressing cells, we employed CLEM. Cells were transfected with the analogous expression construct encoding in addition the NeonGreen gene to allow visualization of transfected cells by fluorescence microscopy (fig. S8D). NeonGreen positive cells were recorded and examined by transmission electron microscopy, revealing abundant clusters of DMVs (fig S8D). Comparison of DMVs induced by nsp3-4 expression and by SARS-CoV-2 infection revealed similar morphology, although expression-induced DMVs were smaller (~125 nm compared to ~300 nm, respectively) (fig S8E). These results show that the sole expression of SARS-CoV-2 nsp3-4 is sufficient to induce DMVs with structural similarity to those generated in infected cells.

Next, we employed this expression-based system to determine AGPAT function in SARS-CoV-2 nsp3-4 induced DMV formation. Huh7-derived cells expressing GFP-tagged AGPAT1 or 2 were transiently transfected with the SARS-CoV-2 HA-nsp3-4-V5 encoding plasmid or the empty vector and colocalization of AGPATs with HA-nsp3 was determined by immunofluorescence microscopy. While in empty vector-transfected cells AGPAT2 and 1 were homogeneously distributed throughout the ER (Fig. 3A and fig. S9A, respectively), we observed a strong relocalization of AGPATs in HA-nsp3-4-V5 expressing cells with AGPATs forming puncta that colocalized with HA-nsp3 (Fig. 3, A and B; fig. S9A). Of note, the relocalization of AGPATs induced by HA-nsp3-4-V5 was not the result of the massive ER alterations occurring in SARS-CoV-2 infected cells, since the subcellular distribution of other ER resident proteins, such as protein disulfide-isomerase (PDI) and calnexin remained unaffected compared to the large puncta observed with AGPATs (Fig. 3C). Since SARS-CoV-2 replication organelles are comprised of DMVs, convoluted membranes and zippered ER ^31^, we next investigated the membrane structures at the sites of AGPAT colocalization with HA-nsp3-4-V5. Using correlative light electron microscopy, we found that relocalized AGPAT puncta perfectly correlated with extensive networks of SARS-CoV-2 HA-nsp3-4-V5 induced DMVs (Fig. 3D). Overall, the data shown here suggest that similar to HCV, AGPATs are relocalized to SARS-CoV-2 nsp3-4 induced DMVs, the likely sites of viral RNA replication ^32^.

**Fig. 3.**
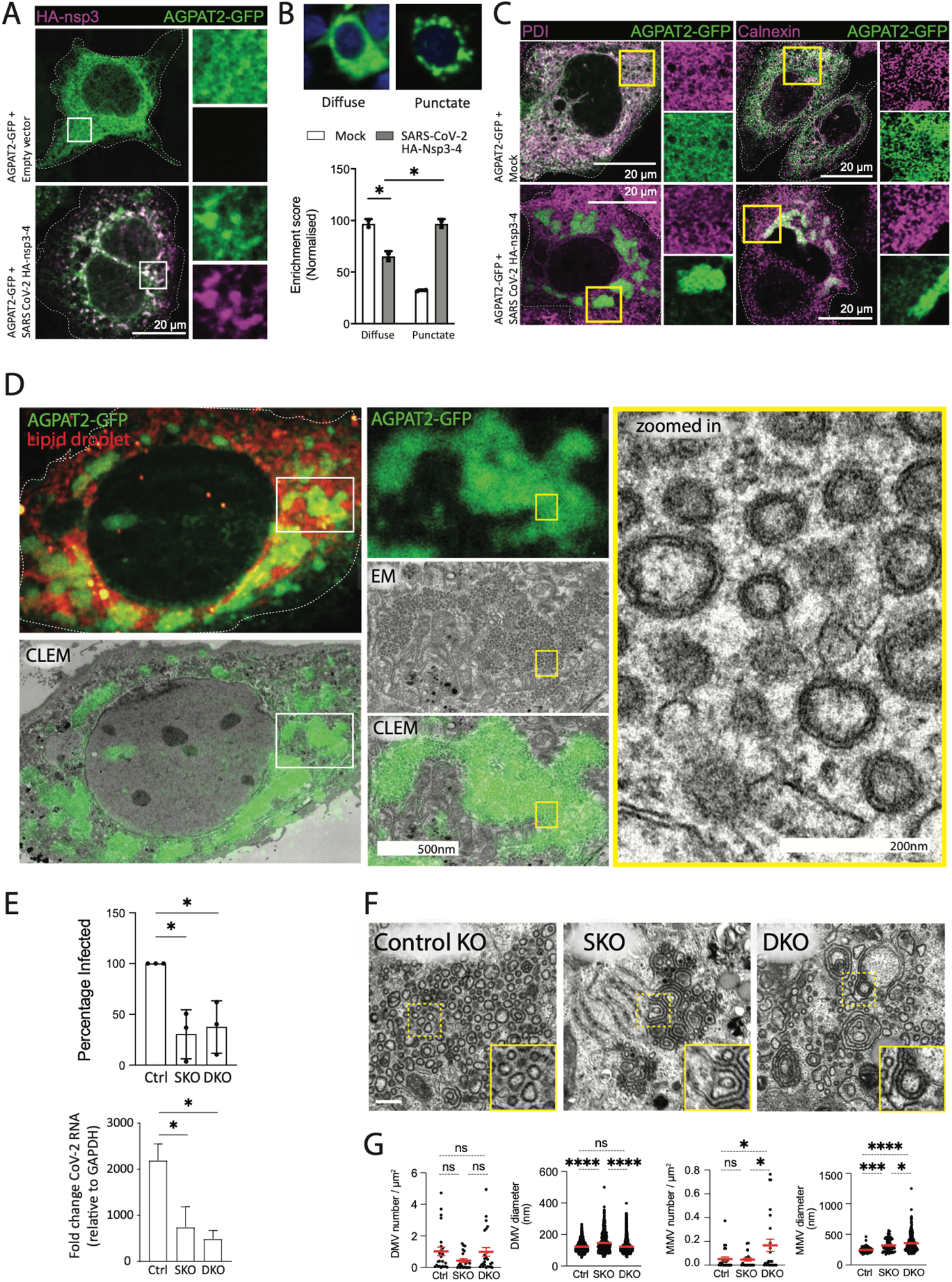
AGPATs are recruited to SARS-CoV-2 induced DMVs and contribute to viral replication. (A) Change of subcellular localization of AGPATs upon expression of SARS-CoV-2 nsp3-4. Huh7-derived cells transiently expressing AGPAT2-GFP were transfected with a SARS-CoV-2 HA-nsp3-4-V5 expression construct or the empty vector. After 48h, cells were stained with HA-specific antibody and examined by confocal microscopy. Maximum intensity projections are shown. Enrichment score indicates the likelihood of cells showing a punctate or diffuse staining pattern. (**B**) Clustering of AGPAT2-GFP in SARS-CoV-2 HA-nsp3-4-V5 expressing cells. Huh7-Lunet/T7 cells were co-transfected with AGPAT2-GFP and SARS-CoV-2 HA-nsp3-4-V5 or the empty vector. Twenty-four hours later, cells were fixed and ~1000 cells per condition were separated into two morphotypes (diffuse or punctate) using CellProfiler Analyst based semi-supervised classifier. Significance was calculated using an unpaired t-test. *, p<0.05. (**C**) AGPAT clustering occurs independent of ER remodeling induced by nsp3-4. Huh7-Lunet cells expressing AGPAT2-GFP and HA-nsp3-4-V5 were stained for the ER markers protein disulfide isomerase (PDI) and calnexin and analyzed by confocal microscopy. (**D**) AGPATs are localized at SARS-CoV-2 HA-nsp3-4-V5 induced DMVs. Huh7-derived cells were transiently transfected with AGPAT2-GFP HA-nsp3-4-V5 and subjected to CLEM. Light and EM images were correlated by using lipid droplets as fiducial markers. White and yellow boxes indicate areas magnified in the corresponding panels on the right. (**E**) AGPAT1/2 contribute to SARS-CoV-2 replication. Huh7-Lunet control, AGPAT2 single (SKO) and AGPAT1/2 double (D)KO cells were infected with SARS-CoV-2 (MOI=0.1). Twenty-four hours later, cells were fixed and immunostained for nucleocapsid, and the percentage of N-positive cells was determined using CellProfiler. Normalized data from three biologically independent experiments are plotted (top right panel). Total RNA was isolated from infected cells, and SARS-CoV-2 RNA levels were determined using RT-qPCR. Data were normalized to cellular GAPDH mRNA (bottom right panel). Significance was calculated using ordinary one-way ANOVA. *, p<0.05. (**F**) Aberrant SARS-CoV-2 DMVs in AGPAT1/2 DKO cells. Huh7-Lunet cells with single (SKO) or double knock-out (DKO) and stably expressing T7 polymerase were transfected with a plasmid encoding SARS-CoV-2 HA-nsp3-4-V5 and fluorescent neon-green. Twenty-four hours later, cells were fixed and NeonGreen positive cells were recorded and examined by EM. HA-nsp3-4-V5 induced DMVs and multi-membrane vesicles (MMVs) were quantified. Shown are the number and diameter of DMVs and MMVs in these cells as observed from at least 8 cell profiles per condition. Statistical significance was calculated using ordinary one-way ANOVA. ****, p<0.001. Light microscopy images in panels A to D are maximum intensity projections.

Next, we tested the effect of AGPAT1/2 depletion on SARS-CoV-2 infection and replication. To this end we used DKO Huh7-Lunet/T7 cells that were employed for the imaging analyses described so far and stably introduced the SARS-CoV-2 receptor gene *ACE2*. Viral replication was measured by using an image-based assay that quantifies the number of cells containing detectable amounts of the nucleocapsid (N) protein (fig. S9B). Using this approach, we observed significant reduction of SARS-CoV-2 positive cells in both single and double AGPAT knockout cells (Fig. 3E). Consistently, RT-qPCR revealed similar reduction of viral replication in single and double KO cells (Fig. 3E, lower right panel). To determine if reduced SARS-CoV-2 replication in AGPAT1/2 KO cells might correlate with altered DMV formation, we transiently expressed SARS-CoV-2 HA-nsp3-4-V5 in control, single and double KO cells. The absence of AGPAT 1/2 did not significantly affect the abundance of cleaved viral proteins HA-nsp3 and nsp4-V5 (fig. S8C). EM analysis of control cells revealed HA-nsp3-4-V5 induced membrane alterations, consistent with an earlier report for MERS-CoV and SARS-CoV ^12^ (Fig. 3, F and G). This included zippered ER and DMVs with an average diameter of 145 nm. In contrast to HCV, the number of nsp3-4 induced DMVs did not decrease in AGPAT single and double KO cells (Fig. 3G, left two panels). However, in both cell pools we observed marked accumulations of multi-membrane vesicles (MMVs), indicating the formation of aberrant membrane structures (Fig. 3, F and G).

To test whether similar to AGPAT1/2 relocalization to nsp3-4 induced DMVs, PA is also enriched at those sites we used the GFP-tagged PA sensor derived from Raf1. In Huh7-derived cells expressing SARS-CoV-2 HA-nsp3-4-V5, the functional version of the sensor (GFP-PABD-Raf1-WT) strongly colocalized with HA-nsp3 in distinct puncta, whereas no such puncta were found with the mutant PABD-Raf1, confirming specificity of PA sensor recruitment to HA-nsp3-containing sites (Fig. 4, A and B).

**Fig. 4.**
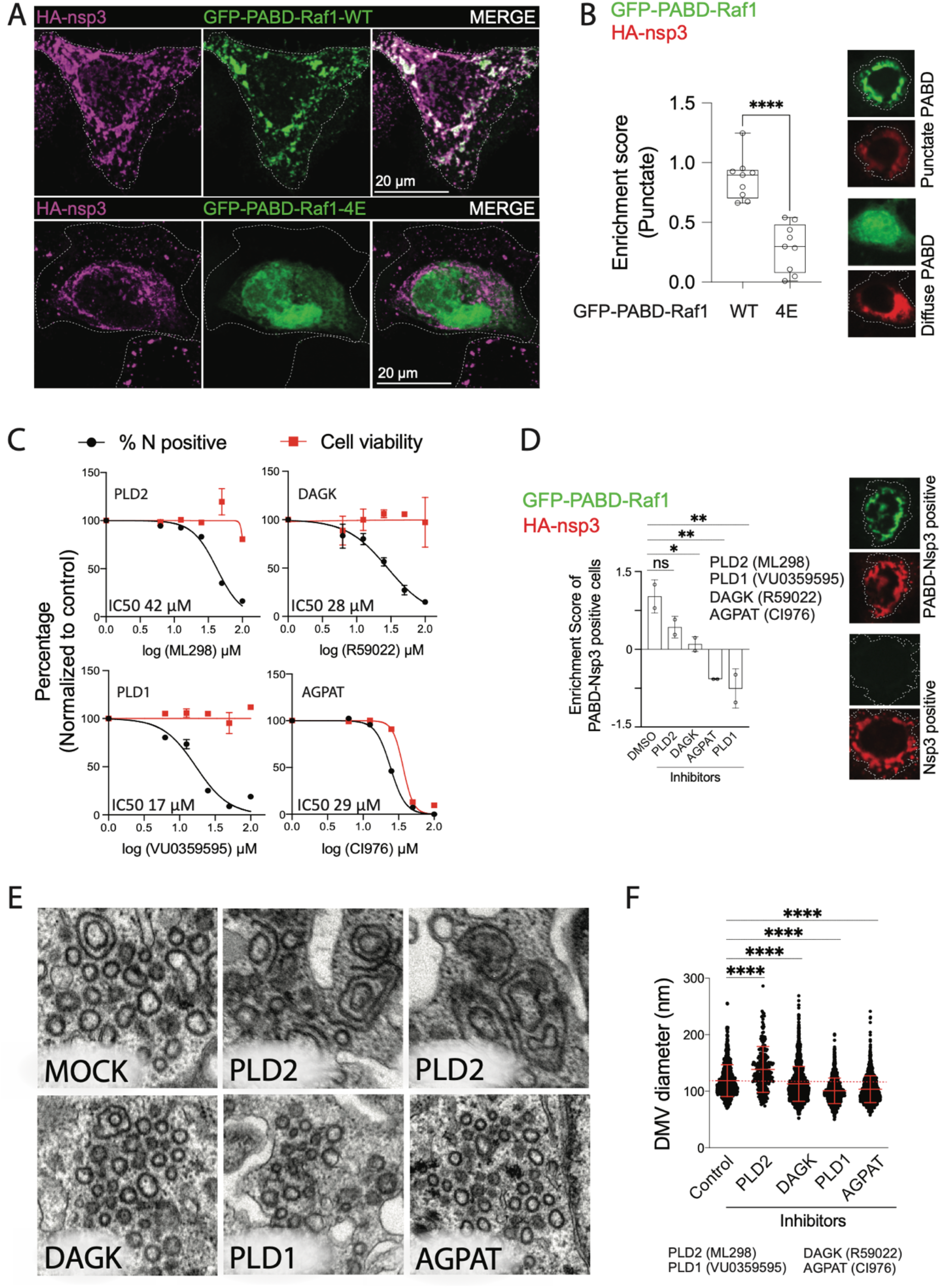
PA accumulation at SARS-CoV-2 DMVs and role of alternative PA synthesis pathways for SARS-CoV-2 replication and DMV formation. (**A**) PA enrichment at SARS-CoV-2 nsp3-containing structures. Huh7-Lunet cells expressing the wildtype or mutant form of the PA sensor were transfected with the plasmid encoding HA-nsp3-4-V5 and 24 h later, cells were fixed, immunostained with HA-specific antibody and HA-nsp3 and GFP-PABD were visualized by confocal microscopy. Maximum intensity projections are shown. (**B**) Using CellProfiler Analyst, a semi-supervised machine learning classifier was trained to differentiate between punctate and diffuse signals of the GFP-PABD sensor (top panel). A normalized enrichment score which indicates the probability of cells showing punctate GFP-PABD localization to nsp3 fluorescent signal across the whole cell population is shown in the graph on the bottom panel. Significance was calculated by unpaired t-test. ****, p<0.0001. (**C**) Alternative pathways for PA generation are important for SARS-CoV-2 replication. Calu-3 cells were infected with SARS-CoV-2 (MOI=5) in the presence of AGPAT, PLD1/2, or DAGK inhibitors. Cells were fixed 24 h post infection, stained with nucleocapsid-specific antibody and percentage of infected cells was quantified using CellProfiler. Cell viability and percentage inhibition are plotted as dose-response curves and IC50 values are given on the top of each panel. (**D**) AGPAT, PLD and DAGK inhibitors reduce PA accumulation at nsp3-positive structures. Huh7-Lunet cells were transfected with SARS-CoV-2 HA-nsp3-4-V5 and GFP-PABD-Raf1 encoding plasmids, followed by addition of a given inhibitor 4h after transfection. Twenty-four hours later, cells were fixed and HA-nsp3 was detected with an HA-specific antibody. GFP-PABD and HA-nsp3 were visualized by confocal microscopy. A semi-automated machine learning based classifier was trained to separate HA-nsp3/PABD double-positive structures from HA-nsp3 single positive structures. Enrichment score for HA-nsp3/PABD double-positive structures showing the up or downregulation of double positive cells in different samples is plotted and statistical significance was calculated using ordinary one-way ANOVA. *, p<0.05, **, p<0.005. (**E**) Decrease of SARS-CoV-2 DMV diameter by AGPAT, PLD and DAGK inhibitors. Huh7-Lunet/T7 cells were transfected with the plasmid encoding HA-nsp3-4-V5 and fluorescent NeonGreen, followed by addition of inhibitors 4 h after transfection. Twenty-four hours later, cells were fixed, NeonGreen positive cells were recorded and examined by EM. Representative images are shown for each condition. (**F**) Number and morphology of DMVs were determined for at least 7 cell profiles per condition. DMV diameters are plotted and statistical significance was calculated using ordinary one-way ANOVA. ****, p<0.001.

Although in comparison to HCV, AGPAT1/2 DKO had lower impact on SARS-CoV-2 replication (compare Fig. 1F with Fig. 3E), and caused a morphologically distinct phenotype of nsp3-4 induced DMVs (Fig. 2A and 3F, respectively), AGPATs, and most likely PA, still accumulated at sites of SARS-CoV-2 DMV clusters (Fig. 4, A and B). This indicates that PA synthesis pathways other than via AGPAT1/2, might contribute to SARS-CoV-2 replication and DMV formation. By means of pharmacological inhibitors of enzymes that convert LPA, PC and DAG to PA (fig. S7A), we measured the dose-dependent effect of these drugs on SARS-CoV-2 replication. All inhibitors reduced SARS-CoV-2 replication in Calu-3 cells and in A549 cells stably expressing ACE2, two commonly used cell models for this virus, at non-cytotoxic concentrations, although in the case of the general AGPAT inhibitor CI976 selectivity was rather low (Fig. 4C and fig. S10A, respectively). Of note, combining the inhibitors at concentrations close to or below their IC50 values caused much stronger reduction of virus replication with no or minimal effect on cell viability, indicating that SARS-CoV-2 can utilize PA produced by alternative PA synthesis pathways (fig. S10, A and B). We then measured the effect of these drugs on PA accumulation at HA-nsp3 containing puncta in HA-nsp3-4-V5 expressing cells and found that all inhibitors reduced PA levels at these sites (Fig. 4D). This reduction was not the result of altered HA-nsp3-4-V5 expression level or self-cleavage, which were unaffected in inhibitor-treated cells (fig. S10C). Next, we determined if reduced PA levels caused by these inhibitors also affect SARS-CoV-2 nsp3-4 induced DMV formation. In cells treated with AGPAT, PLD1, and DAGK inhibitors DMV diameters were significantly reduced (Fig. 4, E and F). Moreover, PLD2 inhibition promoted the formation of MMVs and larger DMVs, similar to what we found in AGPAT single and double KO cells (Fig. 3F). Taken together, our data suggest that PA enrichment is important for proper SARS-CoV-2 DMV formation and viral replication.

Here, we show that PA produced by AGPAT1 and 2 is important for the replication of evolutionary distant positive-strand RNA viruses, HCV and SARS-CoV-2 that amplify their genome in association with DMVs. The remarkable dependence on a common host lipid for the DMV biogenesis in these two viruses that differ profoundly in the diseases they cause and in their biological properties, indicates a striking similarity in the biogenesis of these organelles. Conversely, for viruses replicating their RNA genome in ER-derived membrane invaginations such as the flaviviruses DENV and ZIKV, this lipid pathway appears to be dispensable ^4, 33^. Of note, PA production through AGPAT1 and 2 is also involved in the formation of autophagosome-like DMVs, arguing for some similarity between cellular and viral DMV formation and lipid composition. Additionally, alternative routes of PA biosynthesis contribute to HCV and SARS-CoV-2 replication and DMV generation.

At least three possibilities can be envisioned how PA promotes DMV formation in viral replication and in the context of autophagy. First, the presence of lipids with cone or inverted cone shape in membranes contributes to membrane bending by generating negative or positive membrane curvature, respectively ^16^. While LPA has a large polar head group to fatty acid tail ratio, giving rise to an inverse-cone shape and resulting in positive membrane curvature, the additional fatty acid tail present in PA inverses the head-to-tail ratio. Hence PA displays an overall cone shape, which contributes to negative membrane curvature. Thus, the LPA - PA conversion by AGPATs might contribute to DMV formation by facilitating membrane curvature. Second, PA is directly or indirectly implicated in membrane fission ^34^. This might be achieved by recruitment of effector proteins by PA, either through downstream signaling events, or directly by serving as docking site for PA-binding proteins that have amphipathic or hydrophobic surfaces. In this regard, our NS4B-proteome showed the enrichment of three known PA-interacting proteins, namely, Vitronectin, splicing factor-1, and ubiquitin carboxy-terminal hydrolase L1, in the viral DMV fraction (data S1) ^35^. More than 50 different proteins have been reported to interact with PA, including protein kinases, phosphatases, nucleotide-binding proteins and regulators, however, a comprehensive list remains elusive, and their possible role in the formation of DMVs during autophagy or viral RNA replication, if any, remains to be determined ^18^. Third, an additional role of PA for the functionality of viral replication organelles and perhaps also autophagosomes might be in serving as an exchange lipid in a counter-transporter chain. In the case of HCV, we and others identified accumulation of PI4P at DMVs ^7^ and similar findings have been made for membranous structures involved in the early steps of autophagy ^36^. For HCV, it is thought that PI4P recruits lipid transporters such as OSBP that deliver cholesterol into DMV membranes in exchange for PI4P. A similar mechanism might apply for other lipids or the PI4P precursor PI, with PA serving as a possible exchange factor against these other lipids or PI, respectively.

The similar dependency of DMV-type replication organelles on PA, as reported here for HCV and SARS-CoV-2, might offer an attractive starting point for broad-spectrum antivirals targeting a diverse range of positive-strand RNA viruses replicating in such structures. In line with this assumption, an inhibitor of cytosolic phospholipase A_2α_ has been reported to suppress replication and DMV formation of the 229E human coronavirus and to exert antiviral activity also against the alphavirus Semliki forest virus ^37^. In addition, several human diseases have been linked to defects in PA metabolism and selective autophagy, including neurological disorders and chronic obstructive pulmonary disease ^18, 19^. Although the precise role of PA in these diseases remains to be determined, the critical role of PA for HCV and SARS-CoV-2 infection reported here might offer new approaches for therapeutic intervention.

## Acknowledgements

We thank Marie Bartenschlager, Lena Werstein, Ulrike Herian, Stephanie Kallis, Iris Leibrecht and Fredy Huschmand for excellent technical assistance. We are grateful to Eliana Acosta and Heeyoung Kim for editorial assistance. We acknowledge Alessia Ruggieri for providing empty plasmid constructs. We also acknowledge the excellent support provided by the Infectious Diseases Imaging Platform (IDIP) headed by Vibor Laketa, the University of Heidelberg Electron Microscopy Core Facility (EMCF Heidelberg) headed by Stefan Hillmer and the Proteomics and Metabolomics Facility (Pro-Met-) at CeMM. We thank the European Virus Archive (EVAg) for the provision of the HP/F/2013 ZIKV strain. We also thank all members of the Molecular Virology unit for continuous stimulating discussions.

## Contributions

Conceptualization, K.T., V.P., D.P., and R.B.; Investigation, K.T., V.P., D.P., J-Y.L., M-T.P., W-I.T, C.J.N., M.C., B.C., C-S.T., C.L., P.V., K.H., A.C.M, C.Z., U.H., J.B., L.K., B.B.; Writing – Original Draft, K.T., V.P., and R.B.; Writing – Review & Editing, K.T.,V.P., D.P., J.Y.L., C.J.N., A.M, A.C.M., L.K., H.E., V.T., G.S-F., B.B. and R.B.; Funding Acquisition, R.B., B.B., L.K. and G.S.-F.

## Data and materials availability

All data is available in the main text or the supplementary materials.

## Supplementary Materials

Materials and Methods

Figures S1-S10

Tables S1-S5

References (*1-30*)

Other Supplementary Materials (Data S1-S3)

