## Supplementary Information for "Convergent use of phosphatidic acid for Hepatitis C virus and SARS-CoV-2 replication organelle formation"

**This PDF file includes:**

Materials and Methods

Figs. S1 to S10

Tables S1 to S5

Captions for Data S1 to S3

References

**Other Supplementary Materials for this manuscript include the following:**

Data S1 to S3:

| Table S1 | Proteome analysis |
| --- | --- |
| Table S2 | SiRNA screening |
| Table S3 | Lipidome analysis |

Materials and Methods

Plasmids

To construct the lentiviral vectors pWPI-EGFP-CT, pWPI-EGFP-NT and pWPI-mCherry-NT, EGFP or mCherry coding sequences were amplified by PCR and inserted by in-fusion reaction into the linearized pWPI vector using the BamHI restriction site. Lentiviral plasmids pWPI-AGPAT1-EGFP and pWPI-AGPAT2-EGFP were constructed by insertion of the human AGPAT1 gene (gene ID: 10554) or human AGPAT2 gene (gene ID: 10555) into the BamHI site of the pWPI-EGFP-CT vector.

To construct the expression vector pGEX-PABD and pGEX-PABD_4E, the PA-binding domain (PABD) of the yeast Spo20 gene (gene ID: 855031; nucleotide sequence of the PABD: gacaattgttcaggaagcagaagacgtgataggctacatgtgaagcttaaatccttgaggaataaaatccacaaacaacttcacccaaactgtcggttcgatgacgccactaagactagt) was used. The 4E mutant PABD contained K66E, K68E, R71E, and K73E substitutions as described previously ^1^. PABD_WT or PABD 4E mutant sequences were amplified by PCR and inserted into the pGEX-6P-1 vector using BamHI and XhoI restriction sites. The PABD of Raf1 (gene ID: 5894), which corresponds to amino acid residue 390-426 of the Raf1 protein, was amplified by PCR using the Addgene plasmid 116785 as the source for the DNA. The amplicon was inserted into the pWPI-EGFP-NT vector using HiFi assembly of DNA fragments. The 4E Raf1 PABD mutant contains the following amino acid substitutions as reported earlier: R391E, R398E, K399E and R401E ^2^. The human Parkin gene (gene ID: 5071) was amplified by PCR and inserted into the pWPI-mCherry-NT vector to obtain pWPI-mCherry-Parkin. The PABD T7 expression constructs were constructed using the same approach and the sequences were inserted in pTM1-2eGFP vector to obtain pTM-PABD-Raf1-WT and pTM-PABD-Raf1-4E constructs. All plasmids used in this study are listed in Table S1.

Reagents and resources

All reagents and resources as well as antibodies used in this study are listed in Table S2 and S3, respectively. Phorbol 12-myristate 13-acetate (PMA), Bafilomycin A1 (BafA1), Valinomycin (Val) and Leu-Leu methyl ester hydrobromide (LLOMe) were dissolved in DMSO to prepare stock solutions.

Cell culture and transfection

All cell lines used in this study are listed in Table S4. Cells were maintained in Dulbecco’s modified Eagle medium (DMEM) (Thermo Fisher Scientific), supplemented with 2 mM L-glutamine, nonessential amino acids, 100 U/ml penicillin, 100 μg/ml streptomycin, and 10% fetal calf serum (DMEM cplt). To select for transduced cells, they were cultured in medium containing antibiotics as specified in Table S4. To trigger and visualize starvation-induced autophagy, cells were incubated with serum and amino acid deprived DMEM with or without 200 nM BafA1 for 3 h at 37°C. To monitor mitophagy events, cells were treated with medium containing 10 µM valinomycin at 37°C. To induce PA redistribution, cells were incubated with 100 nM PMA for 5 min. For DNA transfection we used TransIT-LT1 Transfection Reagent according to the manufacturer’s protocol (Mirus Bio LLC). For RNA electroporation, 4x10^6^ cells were suspended in 400 µl cytomix containing 5 µg in vitro transcript, 5 mM glutathione and 2 mM ATP. Electroporation was performed using a Gene Pulser system (Bio-Rad) at 975 µF and 270 V in a 0.4-cm cuvette (Bio-Rad).

Lentivirus production and transduction of cells

Lentivirus production and cell transductions were performed exactly as described earlier ^3^. In brief, HEK-293T cells were co-transfected with the packaging plasmid pCMV-dR8.91, the envelope-encoding plasmid pMD2.G and a pWPI vector plasmid containing the gene-of-interest by use of polyethylenimine (Polysciences Inc.). Supernatants were harvested 48 h and 72 h post-transfection, filtered and virus titers were determined by colony formation assay.

HCV production and viral infection

Production of HCV stocks and infection of cells were performed as described recently ^4^. In brief, Huh7.5 cells were transfected with in vitro transcripts of the HCV variant Jc1 or a renilla-luciferase encoding variant thereof (JcR2A) by electroporation. After 24 h, supernatants were replaced with fresh medium and 48 h and 72 h post-electroporation, supernatants were collected and filtered through 0.45-μm-pore-size filters. Supernatants were stored at −70°C prior to determination of virus titers by limiting dilution assay on Huh7.5 cells. Infected cells were detected by immunohistochemistry using the NS5A-specific antibody 9E10, and the 50% tissue culture infective dose (TCID50) was determined.

Dengue virus and Zika virus production

The reporter virus genomes of DENV-2 (strain 16681s; DV-R2A) and ZIKV (strain H/PF/2013; synZIKV-R2A), which encode renilla luciferase, have been previously described ^5, 6^. DENV stocks were prepared by electroporation of BHK-21 cells with DV-R2A in vitro transcripts and amplified in VeroE6 cells. SynZIKV-R2A stocks were prepared by electroporation of VeroE6 cells or insect C6/36 cells. Extracellular virus titers were determined by plaque forming unit (PFU) assay in VeroE6 cells using an overlay medium containing 1.5% carboxymethylcellulose.

SARS-CoV-2 production

The SARS-CoV-2 isolate Bavpat1/2020 was knidly provided by Prof. Christian Drosten (Charité Berlin, Germany) through the European Virology Archive (Ref-SKU: 026V-03883) at passage 2. Virus stocks were produced in VeroE6 cells by passsaging the virus two times.

Generation of knockout cell lines

The 20-base pairs long guide strands used to target AGPATs are listed in Table S5. CRISPR plasmids were constructed by insertion of annealed oligonucleotides into the lentiCRISPRv2 plasmid (Addgene) encoding a puromycin resistance gene in the case of AGPAT1 or into the lentiCRISPR plasmid (Addgene) encoding a blasticidin resistance gene in the case of AGPAT2. To generate knockout cell lines, cells were transduced with a given lentivirus and two days later cells were cultured in medium containing 3 µg/ml puromycin or blasticidin for at least 3 days. Knock-out was validated by Western blot.

Purification of NS4B-associated membranes and lipid analysis

Pull-down of NS4B-associated membrane fractions was performed as described previously ^7^. In brief, Huh7-Lunet cells (~2.5x10^8^) containing the subgenomic replicon sg4B^HA^31R that encodes HA-tagged NS4B and Huh7-Lunet cells stably overexpressing HA-tagged Calnexin (CNX-HA) and control Huh7-Lunet cells were washed twice with PBS, scraped into PBS, and pelleted by centrifugation at 1,400 × g for 2 min at room temperature. Cells were resuspended in 4 ml hypotonic buffer (20 mM Tris [pH 8.0], 1.5 mM MgCl_2_, 10 mM Na-acetate) and incubated on ice for 30 min. Cells were split into two tubes and disrupted by pressuring 25 times through 2-ml syringes fitted with 22-gauge needles. Nuclei and cell debris were removed by centrifugation at 800 × g for 10 min at 4°C. Post-nuclear supernatants were equilibrated to 150 mM NaCl and subjected to HA-affinity purification. Briefly, 100 ul slurry of HA-antibody coated magnetic beads (Thermo Scientific), were prewashed with 1 ml IP buffer (20 mM Tris [pH 8.0], 150 mM NaCl, 1.5 mM MgCl_2_, 10 mM Na-acetate) and incubated with post-nuclear supernatants on a rotator for 2 h at 4°C. Beads were washed 5 times with 1 ml IP buffer. Bound material was eluted with 50 μl 0.1 M glycine [pH 2.5] for 10 min at room temperature and neutralized with 30 ul of 1 M Tris [pH 7.5]. Eluates of split samples were pooled and subjected to negative staining and Western blot for quality control prior to lipidomic analysis.

For lipidome analysis, membranes released from HA-beads were subjected to an acidic Bligh and Dyer extraction using chloroform/methanol/37% HCl (40:80:1, vol:vol:vol) as previously described ^8^. Extractions were performed in the presence of a lipid standard mix containing 25 pmol phosphatidylcholine (13:0/13:0, 14:0/14:0, 20:0/20:0; 21:0/21:0; Avanti Polar Lipids, Alabaster, AL, USA), 25 pmol sphingomyelin (d18:1 with N-acylated 13:0, 17:0, 25:0, semi-synthesized as described in ^8^, 50 pmol D6-cholesterol (Cambridge Isotope Laboratory), 15 pmol phosphatidylinositol (16:0/ 16:0 and 17:0/20:4; Avanti Polar Lipids), 12.5 pmol phosphatidylethanolamine, 12.5 pmol phosphatidylserine and 5 pmol phosphatidylglycerol (all 14:1/14:1, 20:1/20:1, 22:1/22:1, semi-synthesized as described in ^8^, 12.5 pmol diacylglycerol (17:0/17:0, Larodan), 12.5 pmol cholesteryl ester (9:0, 19:0, 24:1, Sigma-Aldrich, St. Louis, MO, USA), 12 pmol triacylglycerol (LM-6000/D5-17:0/17:1/17:1; Avanti Polar Lipids), 2.5 pmol ceramide and glucosylceramide (both d18:1 with N-acylated 15:0, 17:0, 25:0, semi-synthesized as described in ^8^, 2.5 pmol lactosylceramide (d18:1 with N-acylated C12 fatty acid; Avanti Polar Lipids), (21:0/22:6; Avanti Polar Lipids), and 2.5 pmol lyso-phosphatidylcholine (17:1; Avanti Polar Lipids). Lipid extracts were resuspended in 60 µl methanol and samples were analyzed on an AB SCIEX QTRAP 6500+ mass spectrometer (Sciex, Framingham, MA, USA) with chip-based (HD-D ESI Chip; Advion Biosciences, Ithaca, NY, USA) nano-electrospray infusion and ionization via a Triversa Nanomate (Advion Biosciences) as previously described ^8^. Resuspended lipid extracts were diluted 1:10 in 96-well plates (Eppendorf twin tec 96, colorless, Z651400-25A; Sigma-Aldrich, St. Louis, MO, USA) prior to measurement. Lipid classes were analyzed in positive ion mode applying either specific precursor ion (PC, lyso- PC, SM, cholesterol, Cer, HexCer, and Hex2Cer) or neutral loss (PE, PS, PI, PG, and PA) scanning as described in ^8^.

Data evaluation was performed using LipidView (RRID: SCR_017003; Sciex, Framingham, MA, USA) and an in-house-developed software package (ShinyLipids).

Purification of NS4B-associated membranes for proteome analysis

Pull-down of NS4B-associated membrane fractions was performed as described above, but using Huh7-Lunet cells containing the subgenomic replicon sg4B^HA^31R, and cells containing the analogous replicon with non-tagged NS4B and control Huh7-Lunet cells stably overexpressing HA-tagged Calnexin (CNX-HA). Bound proteins were eluted from the beads by using SDS buffer (50 mM HEPES [pH 7.9], 150 mM NaCl, 5 mM EDTA, 2% SDS). Eluates were subjected to filter aided sample preparation by using a 3 kDa molecular weight cutoff filter (VIVACON 500; Sartorius Stedim Biotech GmbH, 37070 Goettingen, Germany) according to the procedure described earlier ^9^. Fifty microliters of sample were directly mixed in the filter unit with 200 μl of freshly prepared 8 M urea in 100 mM Tris-HCl [pH 8.5] (UA buffer) and centrifuged at 14.000 × g for 15 min at 20°C to remove SDS. Any residual SDS was washed out by two washing steps with 200 μl UA buffer. Proteins were alkylated by incubation with 100 μl 50 mM iodoacetamide in the dark for 30 min at room temperature. After washing three times with 100 μl of UA buffer and three times with 100 µl of 50 mM triethylammonium bicarbonate (TEAB) buffer [pH 8.0] (SIGMA-Aldrich Chemie GmbH, Germany), proteins were digested with 1.25 µg trypsin overnight at 37°C. Peptides were recovered from the filter by centrifugation, applying 40 μl of 50 mM TEAB buffer, followed by 50 μl 0.5 M NaCl (SIGMA-Aldrich Chemie GmbH). Eluted peptides were acidified with TFA, desalted using C18 solid phase extraction spin columns (The Nest Group, Southborough, MA), organic solvent removed in a vacuum concentrator at 45°C and reconstituted in 5% formic acid for analysis by liquid chromatography coupled to tandem mass spectrometry (LC-MS/MS).

LC-MS/MS was performed on a hybrid linear trap quadrupole (LTQ) Orbitrap Velos mass spectrometer (ThermoFisher Scientific, Waltham, MA, USA) coupled to an Agilent 1200 HPLC nanoflow system (Agilent Biotechnologies, Palo Alto, CA, USA) via nanoelectrospray ion source using a liquid junction (Proxeon, Odense, Denmark). Normalized amounts of tryptic peptides (~1 µg) were loaded onto a trap column (Zorbax 300SB-C18 5 μm; 5 × 0.3 mm; Agilent Biotechnologies) at a flow rate of 45 μl/min using 0.1% TFA as loading buffer. After loading, the trap column was switched in-line with a 75 µm inner diameter, 25 cm long analytical column (packed in-house with ReproSil-Pur 120 C18-AQ, 3 μm; Dr. Maisch, Ammerbuch-Entringen, Germany). Mobile-phase A consisted of 0.4% formic acid in water and mobile-phase B of 0.4% formic acid in a mix of 90% acetonitrile and 9.61% water. The flow rate was set to 230 nl/min and a 90 min gradient applied (36 to 30% solvent B within 81 min, 30 to 65% solvent B within 8 min, 65 to 100% solvent B within 1 min, 100% solvent B for 6 min before equilibrating at 36% solvent B for 18 min). For the MS/MS experiment, the LTQ Orbitrap Velos mass spectrometer was operated in data-dependent acquisition (DDA) mode with the 15 most intense precursor ions selected for collision-induced dissociation (CID) in the linear ion trap (LTQ). MS1-scans were acquired in the Orbitrap mass analyzer using a scan range of 350 to 1,800 m/z at a resolution of 60,000 (at 400 m/z). Automatic gain control (AGC) was set to a target of 1×106 and a maximum injection time of 500 ms. MS2-scans were acquired in parallel in the linear ion trap with AGC target settings of 5×104 and a maximum injection time of 50 ms. Precursor isolation width was set to 2 Da and the CID normalized collision energy to 30%. The threshold for selecting precursor ions for MS2 was set to ~2,000 counts. Dynamic exclusion for selected ions was 30 seconds. A single lock mass at m/z 445.120024 was employed ^10^.

Proteome MS data analysis

Acquired raw data files were processed using the Proteome Discoverer 2.2.0.388 platform, utilizing the database search engine Sequest HT. Percolator V3.0 was used to remove false positives with a false discovery rate (FDR) of 1% on peptide and protein level under strict conditions. Precursor masses were recalibrated prior to Sequest HT searches using full tryptic digestion against the human SwissProt database v2017.06 (20,456 sequences and appended known contaminants) with up to one miscleavage site. Oxidation (+15.9949 Da) of methionine and acetylation (+42.010565 Da) of protein N-terminus were set as variable modifications, whilst carbamidomethylation (+57.0214 Da) of cysteine residues was set as fixed modifications. Data was searched with mass tolerances of ±10 ppm and 0.6 Da on the precursor and fragment ions, respectively. Results were filtered to include peptide spectrum matches (PSMs) with Sequest HT cross-correlation factor (Xcorr) scores of ≥1. For calculation of protein intensities, the Minora Feature Detector node and Precursor Ions Quantifier node, both integrated in Thermo Proteome Discoverer were used. Automated chromatographic alignment and feature linking mapping (“matching between run”; https://www.maxquant.org) were enabled. Precursor abundance was calculated using intensity of peptide features including only unique peptide groups.

SAINTexpress version 3.6.3 ^11^ was used to identify interactors in NS4B and CNX pulldowns. The protein intensities obtained from Proteome Discoverer analysis were averaged over technical replicates and used as inputs for the analysis with the SAINTexpress tool. Proteins having SAINT AvgP>0.95 were considered as interactors. All proteins identified by SAINT as interactors (1,543 proteins) of at least one of the baits (NS4B or CNX) were normalized to equal total abundance in each sample analyzed for differential abundance between the NS4B and CNX pulldowns using the limma R software package ^12^. Functional networks for 309 NS4B and 195 CNX significantly enriched proteins (2-fold enriched and having Limma q values of ≤0.05) were generated using ClueGO v2.5.5 app embedded in Cytoscape 3.7.2 ^13^. The human GO (Biological Processes, version from 27 February 2019) was used with the following settings: type of analysis: single; GO terms level: 3–4; GO term restriction: 3 genes and 4%; evidence code: all experimental. A significance threshold level of 0.05 was applied (Data S1).

siRNA screening

siRNA screening was performed by solid-phase reverse transfection of Huh7.5 cells seeded into 96-well plates as described previously ^14^. In brief, siRNA and transfection reagent were seeded into each well of a 96-well plate. After air drying of the plates, 5 x 10^3^ Huh7.5 cells stably expressing firefly luciferase (Fluc) were seeded per well in a volume of 200 µl. After 3 days, cells were infected with the renilla luciferase (Rluc) HCV reporter virus JcR2A ^14^. To determine the impact of knockdown on HCV entry and replication, cells were lysed 72 h after infection and renilla luciferase activity was measured. To account for potential cytotoxic effects of siRNAs, FLuc was measured in the same lysate by using dual luciferase assay. Statistical analysis of the siRNA screening data was performed in R version 3.4, using the Bioconductor package RNAither ^15^. In brief, data were quality-checked, Rluc values were then log-transformed and Lowess normalized against FLuc measurements to account for cytotoxic effects. Measurements were then normalized using z-score normalization with regard to the negative controls, and replicates summarized using the mean.

Luciferase reporter assay

Cells were lysed in 200 µl luciferase lysis buffer (1% Triton X-100, 25 mM glycyl glycin [pH 7.8], 15 mM MgSO_4_, 4 mM EGTA, 10% glycerol) per well in a 12-well plate. Plates were stored at −20 °C until measurement of luciferase activity. For firefly luciferase assay, 200 µl luciferase assay buffer (15 mM K_3_PO_4_ [pH 7.8], 25 mM glycylglycine, 15 mM MgSO_4_, 4 mM EGTA) with freshly added 1 mM DTT, 2 mM ATP and 1 mM D-luciferin (PJK) was mixed with 20 µl luciferase lysis buffer and measured for 20 sec. For renilla luciferase assay, 100 µl luciferase assay buffer supplemented with 1.43 µM coelenterazine (PJK) was mixed with 20 µl lysate and measured for 10 sec by using either a Lumat LB9507 tube luminometer or a Mithras LB940 plate luminometer (both from Berthold Technologies).

Immunofluorescence microscopy

Immunofluorescence microcopy was performed as described previously ^16^. Cells cultured on glass coverslips were fixed with 4% paraformaldehyde in PBS for 30 min. The cells were permeabilized with PBS containing 0.1% Triton X-100, blocked with 5% FBS or BSA, and then incubated with diluted primary antibody for 60 min at room temperature. After washing with PBS three times, cells were incubated with Alexa-dye labeled secondary antibodies in PBS containing 5% FBS for 60 min. The coverslips were mounted in Fluoromount-G (SouthernBiotech) and images were obtained with a Leica SP8 confocal microscope.

Image-based detection of SARS-CoV-2 infection and co-expressing cells

Image-based quantification of SARS-CoV-2 infected cells was based on the immunofluorescence detection of nucleocapsid protein-positive cells. Cells were seeded into 96-well black wall imaging plates for 24 h, followed by infection with SARS-CoV-2 at an MOI of 0.5 in the case of Huh7-Lunet/T7-ACE2 cells or a MOI of 5 for Calu-3 cells. Thirty minutes post infection, inhibitors of PLD1/2 were added and cells were kept at 37˚C for 24 h. The cells were fixed with 4% paraformaldehyde, blocked with 1% skimmed milk, and incubated with anti-nucleocapsid antibody for 1 h at 4˚C, followed by counterstaining with donkey anti-mouse secondary antibody coupled to Alexa568. Nuclear DNA was stained with DAPI and cells were examined using a Nikon Ti2 spinning disk microscope equipped with a Plan Apo lambda 20x/0.75 air objective and a back-illuminated EM-CCD camera (Andor iXon DU-888). Segmentation of nuclei were done with the CellProfiler version 3.1.9 software package. Cytoplasm was identified by expanding the nuclei by 5 pixels. Separation of cells into infected and non-infected population was performed with a semi-supervised machine-learning based approach using CellProfiler Analyst, as described earlier ^17^. The schematics of the pipeline is outlined in fig S9B. The enrichment score of cells co-expressing HA-nsp3 and GFP-PABD-Raf1 (in Fig 4D) or expressing punctate pattern of AGPAT2 (Fig. 3B) and GFP-PABD-Raf1 (Fig. 4B), is calculated as described before ^18^. In short, the enrichment score indicates the probability of the presence of a specific class (for example, co-expression or punctate expression pattern) in different samples in relation of the total cells in the samples.

RT-qPCR assay for SARS-CoV-2 replication

Total RNA was extracted from cells using NucleoSpin RNA extraction kit (Machery-Nagel) following manufacturers protocol. Reverse transcription (RT) reaction for cDNA synthesis was performed using the high capacity cDNA RT kit (ThermoScientific). Each cDNA was diluted 1:5 in nuclear free H2O and qPCR was performed using iTaq Universal SYBR green mastermix (Bio-Rad). Primers for qPCR were designed using Primer3 for SARS-CoV-2-ORF1 (Forward 5’- GAGAGCCTTGTCCCTGGTTT-3’, Reverse 5’-AGTCTCCAAAGCCACGTACG-3’) and HPRT (Forward 5’-CCTGGCGTCGTGATTAGTG-3’, Reverse 5’-ACACCCTTTCCAAATCCTCAG-3’). Relative abundance for SARS-CoV-2 Orf1 mRNA was corrected for PCR efficiency and normalized to HPRT transcript level.

SARS-CoV-2 DMV generation and quantification

Based on earlier findings that coronavirus DMV formation can be induced by the sole expression of viral nonstructural protein (nsp)3-4 ^19^, we similarly constructed an expression vector containing the SARS-CoV-2 (nsp)3-4 genes tagged with HA (for nsp3) and V5 (for nsp4). Expression of this construct in Huh-derived cells showed significant number of DMVs in only the transfected cells, independent of infection. Using this system, we measured the number and diameter of DMVs and MMVs in Lunet cells under different perturnation modalities (Fig 3F, 4E and F), as described before for HCV-induced DMV formation ^20^.

Immunoprecipitation and immunoblotting

For immunoprecipitation, cells were processed as described for proteome analysis, but captured protein complexes were eluted by using 2 x sample buffer (100 mM Tris-HCl [pH 6.8], 4% SDS, 12% β-mercaptoethanol, 20% glycerol, 0.001% bromophenol blue). To prepare lysates for immunoblotting, cells were incubated in 2× lysis buffer (200 mM Tris [pH 8.8], 5 mM EDTA, 0.1% bromophenol blue, 10% sucrose, 3% SDS, 2% β-mercaptoethanol) and incubated for 5 min at 95°C. Proteins were separated by SDS-polyacrylamide gel electrophoresis and electro-transferred onto PVDF membranes. After blocking of the membranes with 5% nonfat milk, they were incubated overnight at 4°C with primary antibodies. After washing with 0.5% Tween 20 in PBS, membranes were incubated with secondary horseradish peroxidase-conjugated antibodies for 1 h at room temperature. Membranes were developed by using Western Lightning Plus-ECL reagent (PerkinElmer), and signals were detected by Intas ChemoCam Imager 3.2 (Intas).

Electron microscopy

Transmission EM was performed as described previously ^7^. In brief, cells were fixed with 2.5% glutaraldehyde (GA) in 50 mM sodium cacodylate buffer (CaCo), supplemented with 2% sucrose, 50 mM KCl, 2.6 mM MgCl_2_ and 2.6 mM CaCl_2_, for 30 min at room temperature. After five washes with 50 mM CaCo, samples were incubated with 2% OsO_4_ in 25 mM CaCo for 40 min on ice, washed three times with EM-grade water and incubated in 0.5% uranyl acetate in water overnight at 4°C. Samples were rinsed three times with EM-grade water, dehydrated in a graded ethanol series (from 40% to 100%) at room temperature, embedded in Epon 812 (Electron Microscopy Sciences) and polymerized for 2 days at 60°C. After polymerization, ultrathin sections of 70 nm were obtained by sectioning with an ultramicrotome Leica EM UC6 (Leica Microsystems) and mounted on a slot grid. Sections were counterstained using 3% uranyl acetate in 70% methanol for 5 min and lead citrate (Reynold’s) for 2 min and examined by using a JEOL JEM-1400 (JEOL) operating at 80kV and equipped with a 4K TemCam F416 (Tietz Video and Image Processing Systems GmbH).

Correlative light and electron microscopy (CLEM)

Two methods were employed using protocols described previously ^21^. For CLEM analysis with low resolution fluorescence imaging, 0.5x10^5^ Huh7-Lunet/T7 cells were seeded onto glass-bottom culture dishes containing gridded coverslips (MatTek Corporation) and incubated overnight. Cells were transfected with plasmid pTM NS3-5B encoding the HCV replicase (NS3, NS4A, NS4B, NS5A and NS5B) with NS5A containing a GFP insertion that does not affect viral protein function ^22^. For Fig 3D, plasmid encoding AGPAT2-GFP was transfected into Huh7-Lunet/T7 cells. After 24 h, differential interference contrast (DIC) and GFP signals were acquired by using a widefield fluorescence microscope (Nikon Eclipse) with a 10x objective lens. Cells were fixed with 2.5% GA, 2% sucrose in 50 mM CaCo, supplemented with 50 mM KCl, 2.6 mM MgCl_2_ and 2.6 mM CaCl_2_ for 30 min at room temperature. After five washes with CaCo, cells were processed for EM analysis as described above.

For CLEM with high precision image correlation, cells were fixed with 4% paraformaldehyde and 0.2% GA in PBS for 30 min at room temperature, washed three times with 150 mM glycine in PBS, once with PBS, stained with LipidToxTM Deep Red Neutral Lipid Stain (Invitrogen) and analyzed by using a spinning disc confocal microscope. Fluorescence images were acquired with optical sections of 0.2 µm using a 100x objective. For low-precision CLEM (Fig 3F and 4E), Huh7-Lunet/T7/Ctrl KO, Huh7-Lunet/T7/AGPAT2 KO or Huh7-Lunet/T7/DKO cells were seeded onto dishes containing gridded coverslips (MatTek Corporation). Twenty-four hours after transfection with pTM-SARS-CoV-2-HA-3-4-V5-2A-NG (NeonGreen) using TransIT-LT1 Transfection Reagent (Mirus Bio), samples were observed by confocal microscopy with a 10x objective lens to locate NeonGreen-positive cells. After imaging, cells were fixed again with 2.5% GA, 2% sucrose in 50 mM CaCo, supplemented with 50 mM KCl, 2.6 mM MgCl_2_ and 2.6 mM CaCl_2_ for at least 30 min on ice. Further processing for EM analysis was the same as described above.

Detection of PA in transiently permeabilized cells

A recombinant protein composed of the PA-binding domain (BD) from yeast Spo20 fused to the N-terminus of glutathione-S-transferase (GST-PABD) was expressed in E. coli, strain BL21 (DE3), by using the expression vector pGEX-6P-1. Cells were grown in LB medium until OD600 = 0.5, followed by a 3 h-incubation period in medium containing 1 mM isopropyl β-d-1-thiogalactopyranoside (IPTG) at 37°C. Cells were harvested by centrifugation and GST fusion proteins were purified from cell lysates by using a Spin Purification kit as recommended by the manufacturer (Thermo Fisher Scientific). Eluates were dialyzed in PBS using Slide-A-Lyzer Dialysis Cassettes (7K molecular weight cut-off; Thermo Fisher Scientific) and purified proteins were stored in 50% glycerol in PBS at -20°C at a concentration of 1 mg/ml. Detection of PA with the GST-PABD in semi-intact cells was performed as described previously with slight modifications ^23^. In brief, cells seeded on coverslips were washed twice with PBS and incubated on ice with 200 ng/ml streptolysin O (SAE0089; Sigma Aldrich) in PBS for 5 min. After washing three times with PBS, cells were incubated with transport buffer (25 mM HEPES-KOH, [pH 7.4], 115 mM potassium acetate, 2.5 mM MgCl_2_) at 37 °C for 5 min. After washing twice with transport buffer at room temperature, cells were incubated with a reaction mixture (1 mM ATP, 50 µg/ml creatine kinase, 2.62 mg/ml creatine phosphate, 1 mg/ml glucose, 1 mM GTP, and 5 μg/100 μl GST-PABD wild-type or 4E mutant) at 37°C for 15 min. Cells were washed with transport buffer, fixed with 4% paraformaldehyde for 30 min, and permeabilized with 0.2% Triton X-100 for 20 min. After blocking with 5% skim milk, GST-PABD proteins were detected using a GST-specific mouse monoclonal antibody as primary antibody and an Alexa 488-conjugated anti-mouse antibody as a secondary antibody.

Cell viability and cell growth assays

Cell viability was measured by using the CellTiter-Glo luminescent cell viability assay (Promega) according to the protocol of the manufacturer. Cell growth was determined by cell counting using a TC20 automated cell counter (BioRad).

Statistics and reproducibility

Unless otherwise stated, values represent the mean of a given number of replicates. Error bars are SD or SEM as indicated in the figure legends. Student’s t-tests were performed for unpaired or paired groups by using the GraphPad Prism 5.03 software package (GraphPad software). A *P* value < 0.05 was considered statistically significant. All experiments were repeated two or three times independently, as indicated in the figure legends. No statistical method was used to predetermine sample size. The experiments were not randomized and the investigators were not blinded to allocation during the experiments.

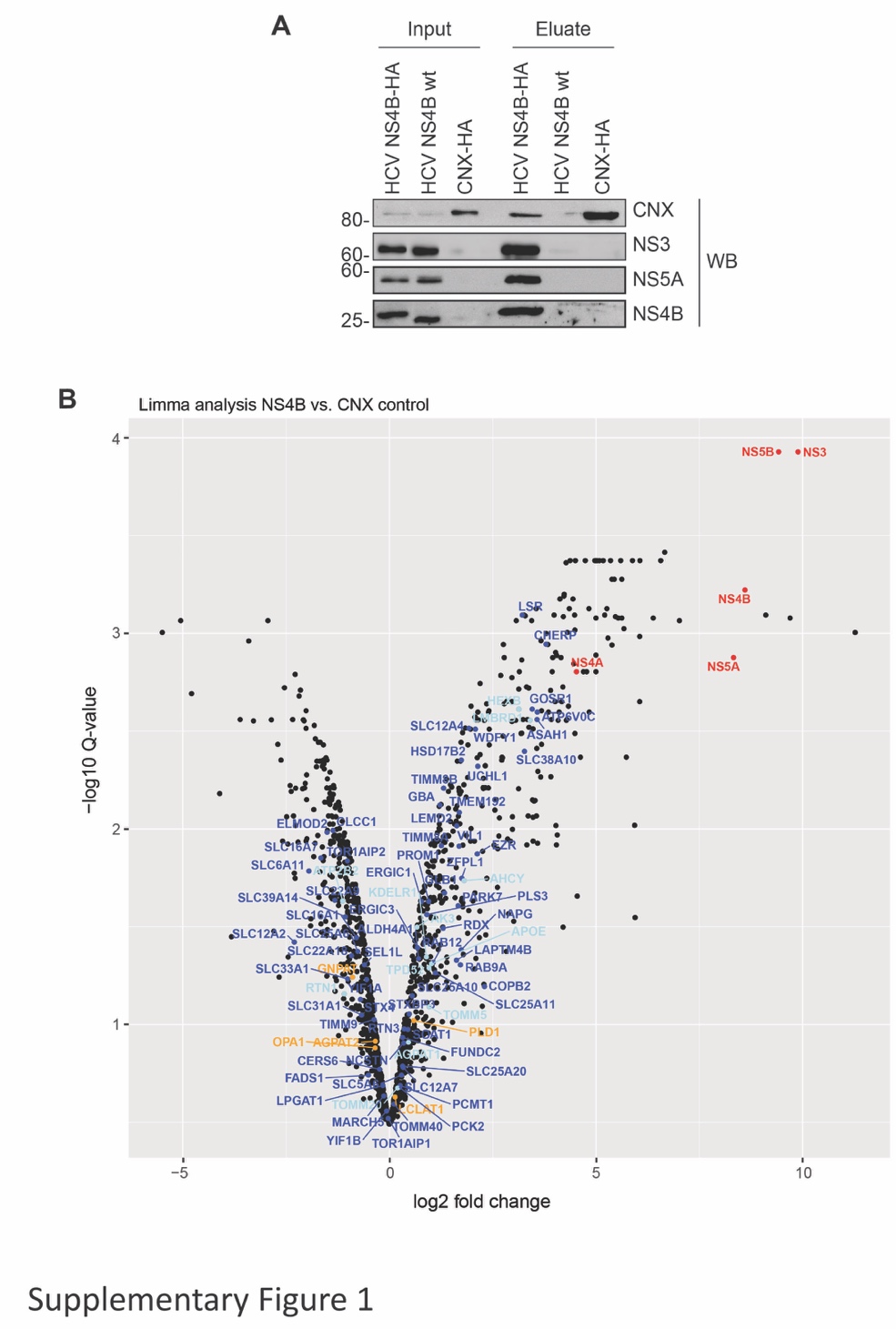

Fig. S1. Purification and proteome of NS4B-associated membranes. (A) NS4B-associated DMV fractions were isolated from Huh7-Lunet cells containing a stable subgenomic HCV replicon of the isolate JFH1. The replicon encoded either wildtype or an HA-tagged NS4B (NS4B-wt and NS4B-HA, respectively). Huh7-Lunet cells stably expressing HA-tagged calnexin (CNX-HA) were used as biological reference. After purification under native conditions, which were used to preserve the membranes, 10% of the eluate and 0.5% of the cell lysate (input) were analyzed by western blotting (WB). The replicon with wildtype NS4B served as technical control. The remainder of the samples was used for mass-spectrometry to determine total protein composition. All three conditions were prepared in duplicates and analyzed by LC-MS/MS based proteomics. (B) Volcano plot of differentially enriched interactors of NS4B and CNX. The q-value was calculated using the limma R software package [2] and corrected for multiple hypothesis testing. Proteins selected for validation are highlighted by dark blue, proteins confirmed by siRNA screen in orange and virus proteins in red color.

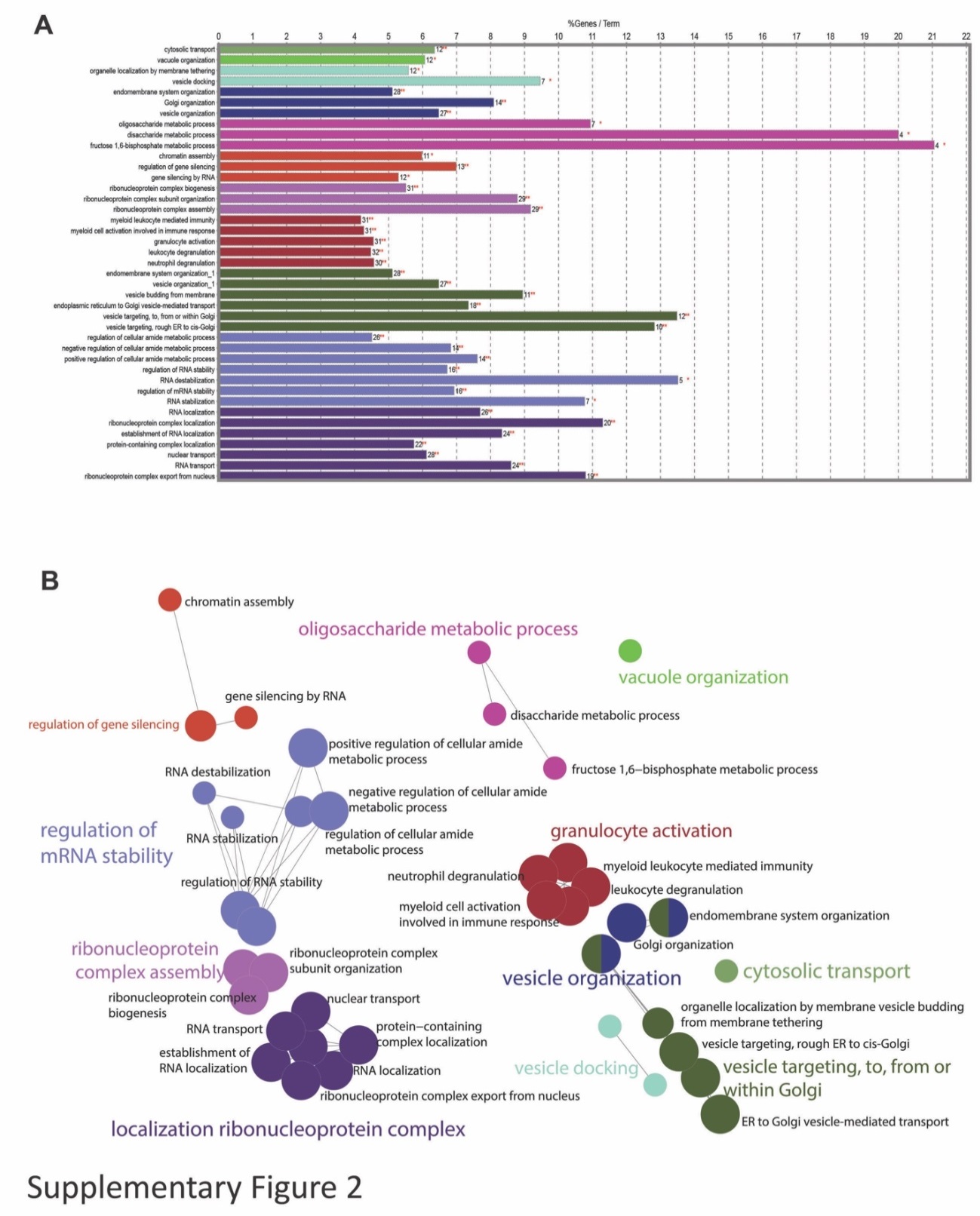

Fig. S2. Gene ontology analysis of proteome hits identified in NS4B-enriched membrane fractions. List of 309 proteins showing a ≥2-fold enrichment in NS4B-associated DMV fraction versus HA-tagged calnexin (CNX-HA) and having limma statistical q-values higher than 0.05 were used for gene ontology (GO) analysis. Cytoscape 3.7.2 ^13^ and ClueGO ^24^ app were used to generate and visualize non-redundant biological terms for large clusters of protein accessions (human uniprot accession IDs) in a functionally grouped network. ClueGO results are illustrated as a functionally grouped network of terms. (A) Bar chart of ClueGO significant biological functional terms organized in multiple term occurrences. Numbers represent protein accessions per term. (B) ClueGO results represented as functionally grouped network of terms with node size showing significance of cluster.

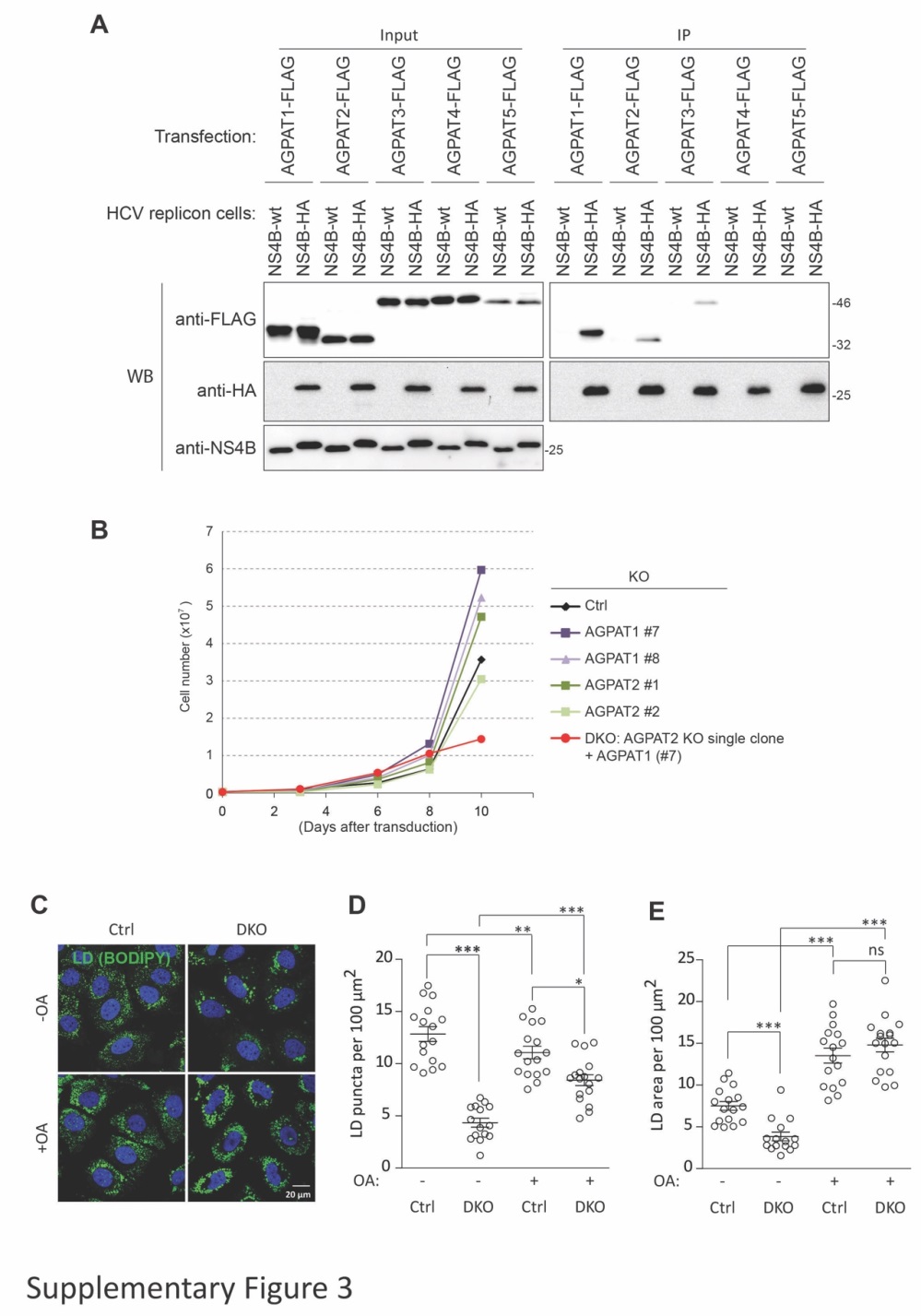

Fig. S3. Interaction of NS4B with AGPAT isoforms and impact of single and double AGPAT1/2 knock-out on cell viability as well as lipid droplet formation. (A) Huh7-Lunet cells containing a stably replicating HCV replicon with wildtype or HA-tagged NS4B were transfected to express FLAG-tagged AGPATs. Two days after transfection NS4B-HA enriched membrane fractions were prepared under native conditions by HA-specific immunoprecipitation (IP). Proteins in lysates (1% of input) and captured protein complexes (20% of eluate) were analyzed by western blotting (WB). (B) Effect of AGPAT1/2 single and double knock-out on cell growth. Cells were infected with sgRNA encoding lentivirus on day-0 and expanded until day-10 using increasingly bigger culture dishes according to cell growth. Cell numbers were determined by using an automated cell counter. The experiment was repeated twice. (C to E) Control KO and AGPAT1/2 DKO cells were incubated without or with 50 µM oleic acid-BSA (-OA and +OA, respectively) for 16 h. (C) Representative images. Lipid droplets (LD) and nuclear DNA were stained with BODIPY 493/503 (green) or DAPI (blue), respectively. (D and E) The number of LD puncta per 100 µm^2^ cell surface area (D) or total LD area per 100 µm2 (E) were determined and are given as average and SEM. Significance was calculated by a paired t-test. *, p<0.05; **, p<0.01; ***, p<0.0001.

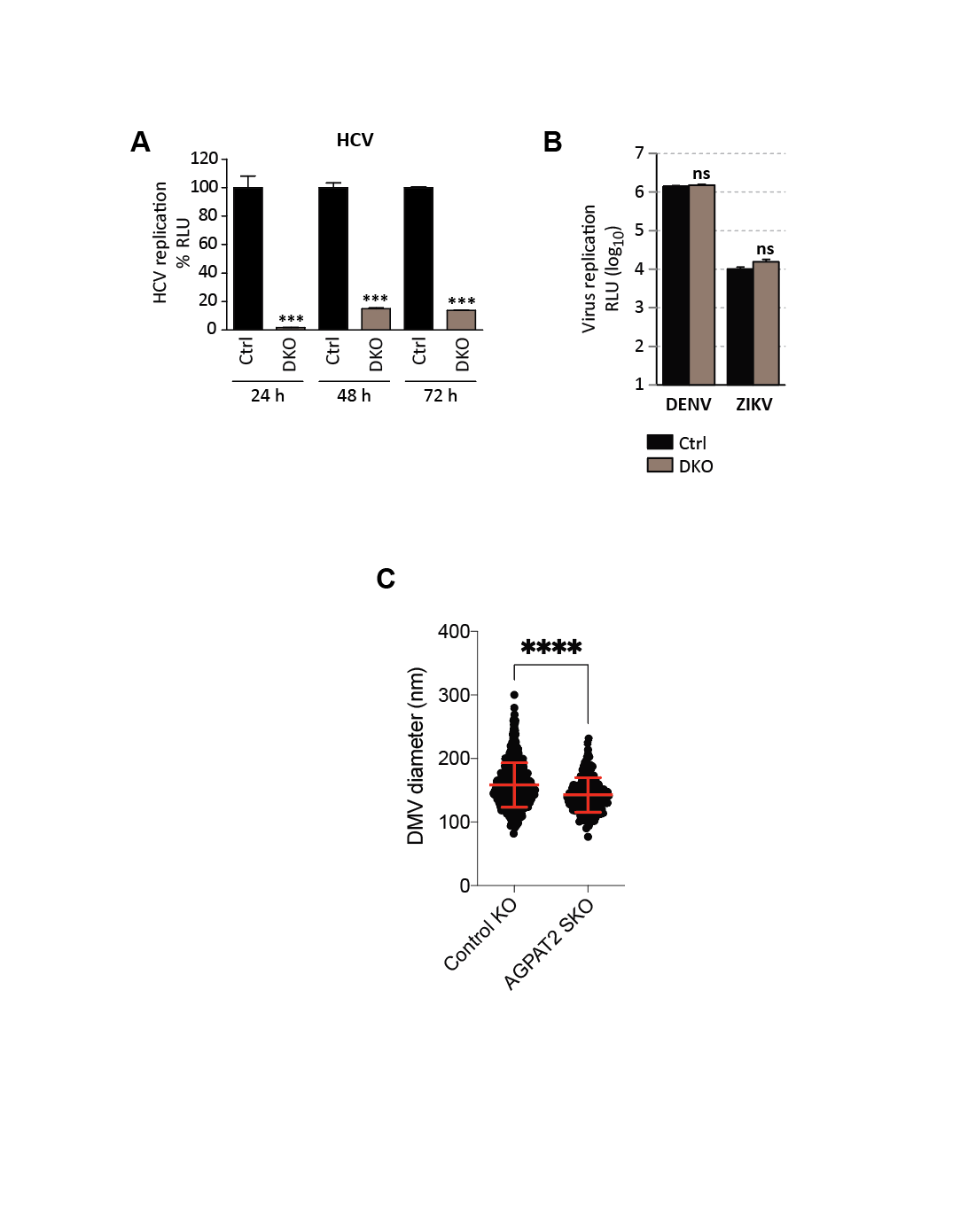

**Fig. S4. Effect of AGPAT1/2 DKO on replication and DMV formation of HCV and on replication of other flaviviruses.** (**A**) Huh7-Lunet/T7 cells were electroporated with in vitro transcripts of a subgenomic HCV reporter replicon encoding the firefly luciferase. Luciferase activities were analyzed at indicated time points after electroporation. Graph shows average and SD from 3 independent experiments. Significance was calculated by a paired t-test. ***, p<0.001. (**B**) AGPAT1/2 DKO does not affect DENV and ZIKV replication. Cells were infected with DENV or ZIKV *renilla* luciferase reporter viruses and 48 h later, RNA replication was determined by luciferase assay. Values are expressed as average of RLU (log_10_) and SD from 3 independent experiments. Significance was calculated by paired t-test. ns, p>0.05. (**C**) Reduced HCV DMV diameter in AGPAT2 single (S)KO cells. Huh7-derived cells stably expressing the T7 RNA polymerase and without or with SKO of AGPAT 2 were transfected with the HCV replicase-encoding plasmid containing a GFP insertion in NS5A (construct pTM NS3-5B/5A-GFP) ^21^. After 24 h, cells were fixed and subjected to CLEM. DMV diameters within whole cell sections were counted and plotted. Significance was calculated by paired t-test. **, p<0.0001.

­
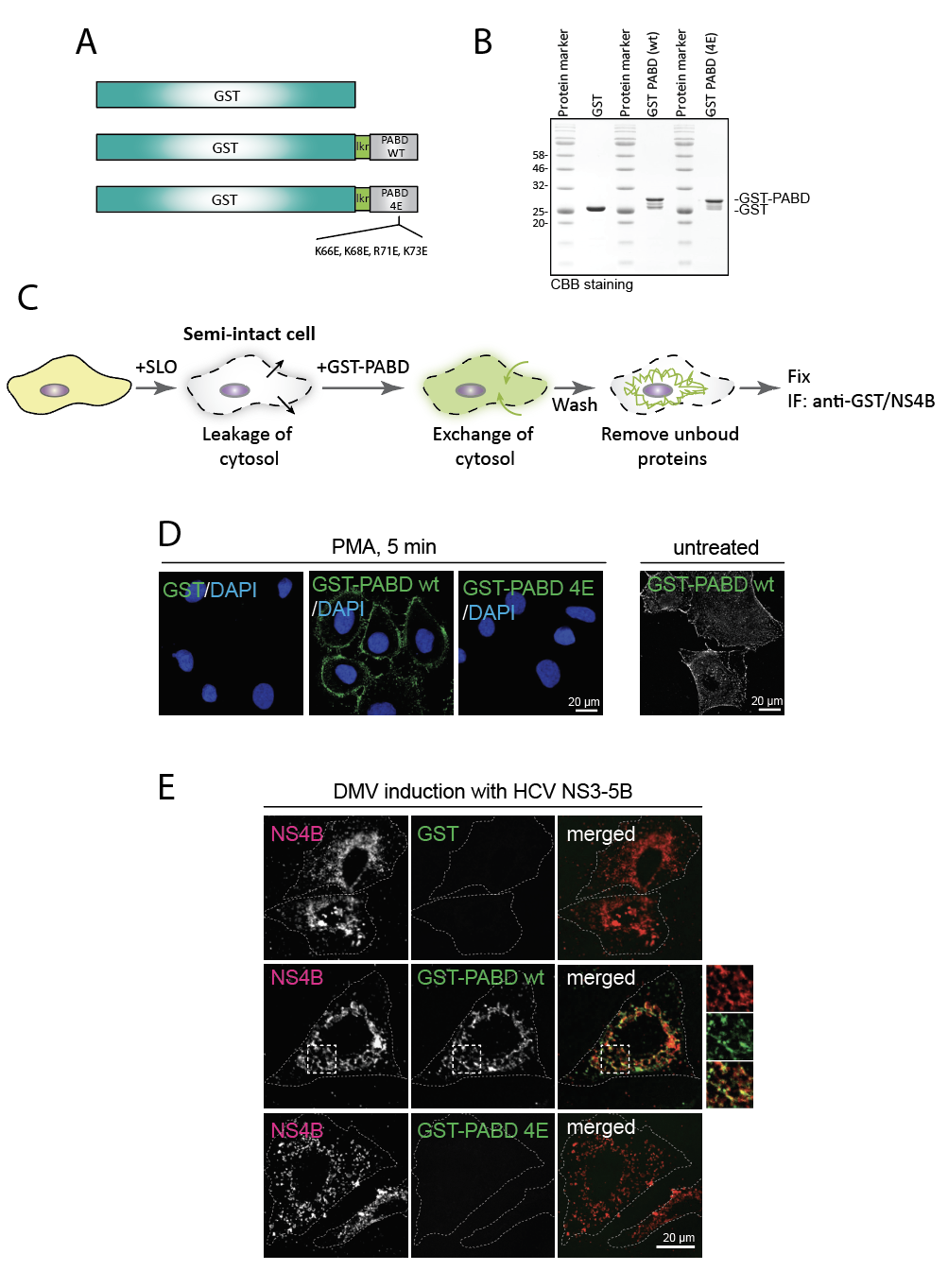

**Fig. S5. Subcellular PA distribution in HCV replicase-expressing cells as determined with a recombinant PA sensor.** (**A**) Schematic of the recombinant PA-binding protein that is composed of the glutathione S-transferase (GST) fused to the PA binding domain (PABD) of the Spo20 protein via a short linker (lkr). The 4E mutant contains 4 amino acid substitutions specified on the bottom. (**B**) Fusion proteins were expressed in E. coli and pre-purified cell lysates were subjected to GST-specific affinity chromatography. Five microgram recombinant protein were loaded onto a SDS-polyacrylamide gel that was stained with coomassie brilliant blue (CBB) after electrophoresis. (**C**) Experimental approach to visualize PA with the exogenously added recombinant biosensor. Cells were transiently permeabilized by treatment with streptolysin O (SLO) that forms pores in the plasma membrane and allows the partial exchange of the cytosol against a physiological solution containing the purified recombinant PA biosensor. Bound GST-PABD and NS4B were visualized by immunofluorescence (IF) microscopy using GST- and NS4B-specific antibodies. (**D**) Functionality of the recombinant PA sensor. Huh7-Lunet/T7 cells were treated with 100 nM PMA for 5 min or left untreated and PA was visualized by IF using recombinant proteins specified on the top of each panel. Nuclear DNA was stained with DAPI to visualize all cells on the coverslip. (**E**) Huh7-Lunet/T7 cells were transfected with the HCV NS3-5B encoding plasmid, followed by permeabilization with SLO and addition of purified GST-PABD wt or 4E proteins. Bound proteins were detected by GST-specific immunofluorescence microscopy.

­­
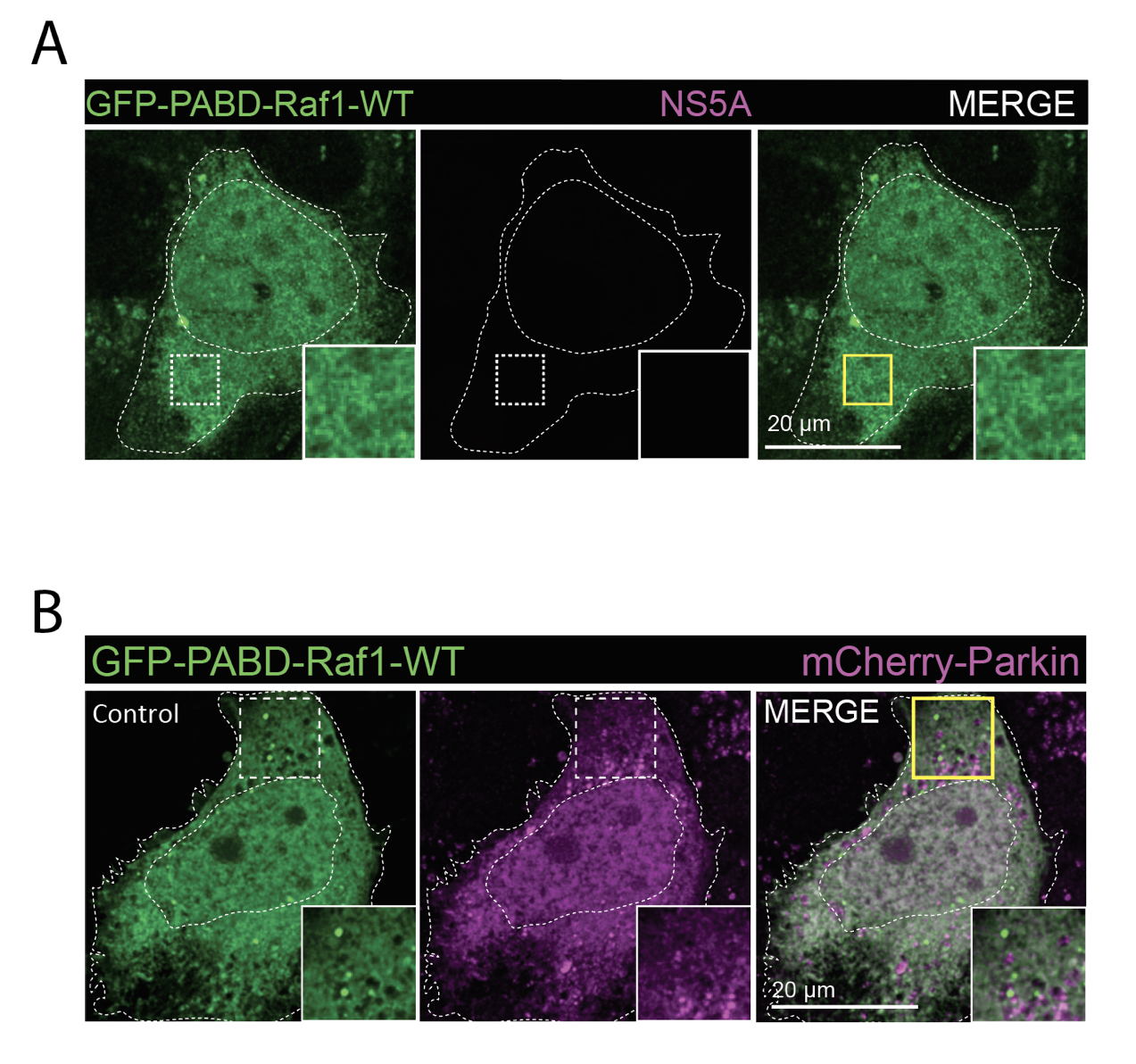

**Fig. S6. PA distribution in naïve Huh7-derived cells. (A)** Huh7-Lunet/T7 cells were transfected with a construct encoding EGFP-tagged wildtype (WT) PA sensor (construct pTM-EGFP-PABD-Raf1-WT). Twenty-four hours later, cells were fixed and GFP-PABD was visualized by fluorescence microscopy. White boxes indicate regions magnified in the lower right of each panel. Data shown here belong to the results shown in Fig. 2E and they reveal a diffuse PA-sensor distribution in the absence of HCV NS3-5B expression. (**B**) Subcellular PA-sensor and mCherry-Parkin distribution in the absence of mitophagy. Huh7-derived cells were co-transfected with EGFP-PABD-Raf-1 and mCherry-tagged Parkin and 24 h later cells were fixed and analyzed by fluorescence microscopy. Data shown here belong to the results shown in Fig. 2G.

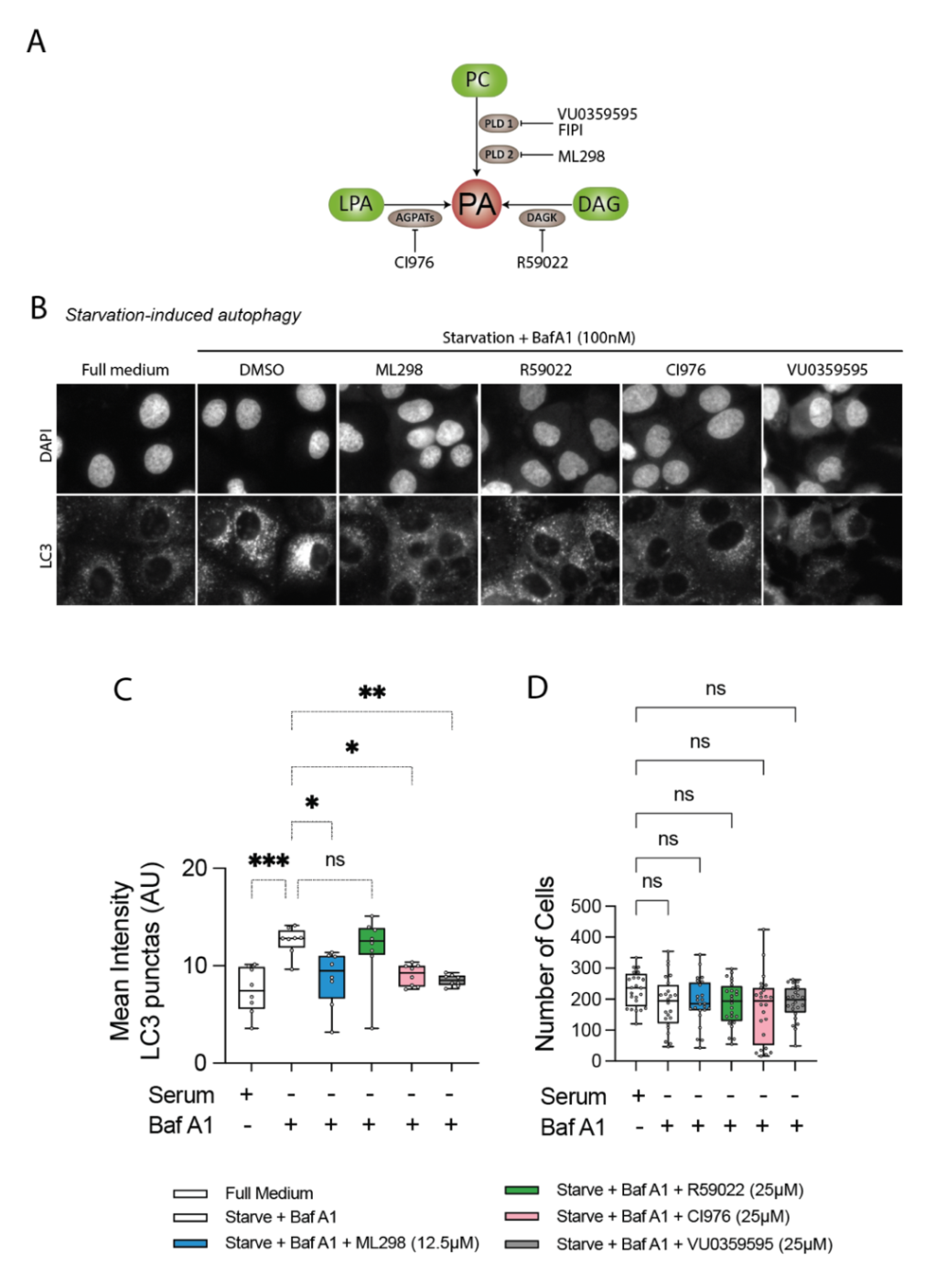

**Fig. S7. Inhibition of PA production through PLD1, PLD2 and AGPATs decreases LC3 accumulation during nonselective autophagy.** (**A**) PA biosynthesis pathways via lysophosphatidic acid (LPA), phosphatidylcholine (PC) and diacylglycerol (DAG), metabolized by AGPATs, PLDs and DAGK, respectively. (**B**) Effect of PA inhibitors on LC3 puncta. Huh7-Lunet/T7 cells were incubated in full medium or starvation medium containing BafA1, in the presence of PA inhibitors specified above the panels. After 3 h incubation, cells were fixed and stained with an LC3-specific antibody followed by immunofluorescence microscopy. Nuclear DNA was stained with DAPI. Solvent (DMSO) treated cells served as control. (**C**) Quantification of mean LC3 puncta intensity per cell, and (**D**) total cells analyzed per condition, was determined by using a custom-made CellProfiler script that segments the LC3 puncta, nuclei and cell boundary and measures the intensities of LC3 puncta in the cytoplasmic area of single cells. Each dot in the scatter plot represents an imaging field of up to 500 cells. Significance was calculated using ordinary one-way ANOVA. *, p<0.05. **, p<0.01. ***, p<0.001, ns, p>0.05.

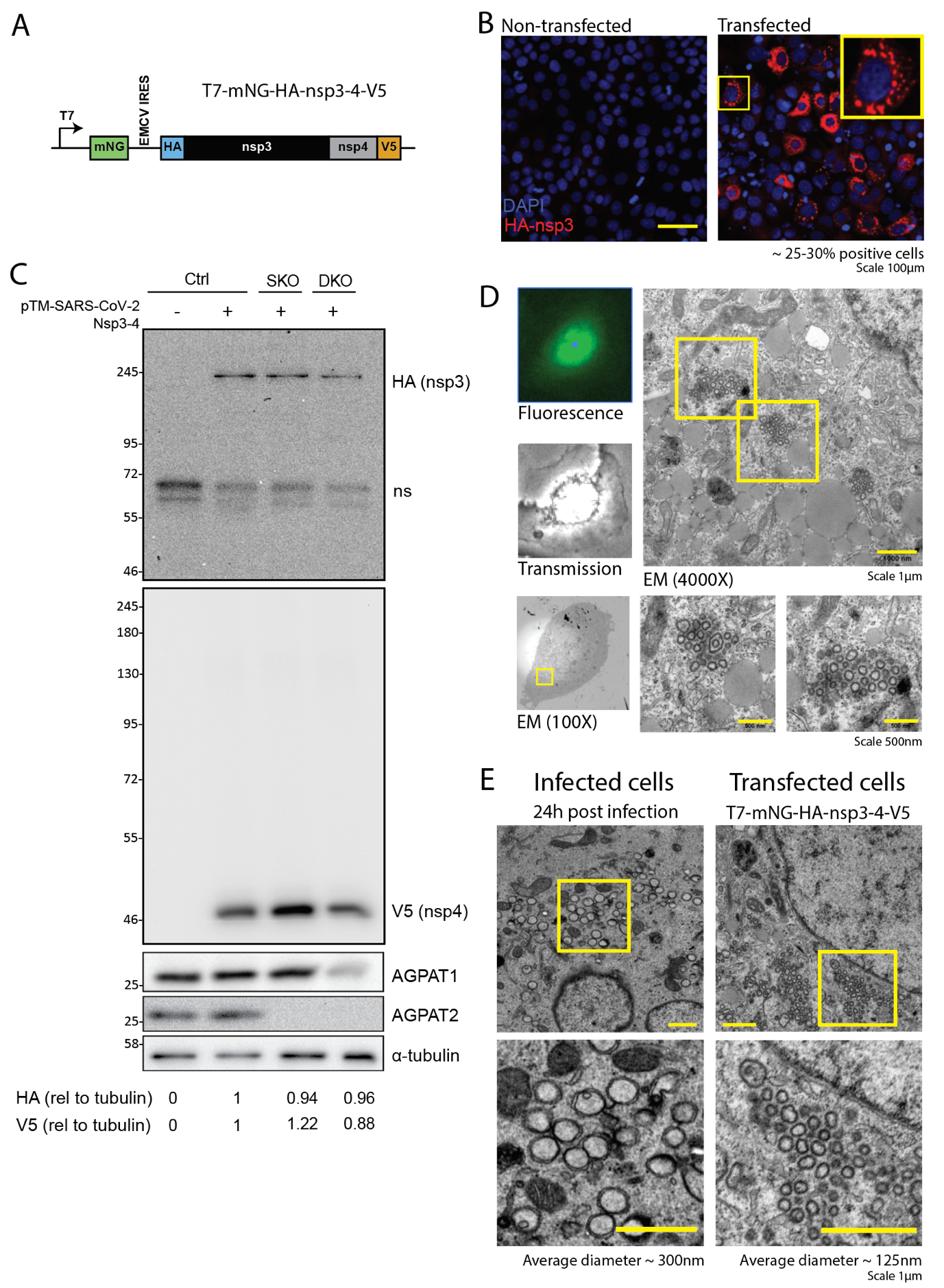

**Fig. S8. SARS-CoV-2 nsp3-4 induced DMVs are morphologically similar to infection induced DMVs, and effect of AGPAT KO on viral protein expression. (A)** Schematics of the T7-based expression construct encoding SARS-CoV-2 HA-nsp3-4-V5. mNG, NeonGreen. **(B)** Huh7-derived cells were transfected with the SARS-CoV-2 HA-nsp3-4-V5 encoding plasmid and 24 h later cells were stained with an HA-specific antibody and analyzed by confocal microscopy to visualize HA-nsp3. Transfection efficiency given on the lower right was determined by cell counting. A magnification of the yellow boxed area is shown on the top right. **(C)** Abundance of SARS-CoV-2 proteins in transfected cells was quantified by western blotting using primary antibodies specified on the right of each panel. α-tubulin served as loading control. Data in **(C)** belong to the results shown in Fig. 3F and G. ns, non-specific band. **(D)** CLEM for quantification of nsp3-4 induced DMVs. Huh-derived cells stably expressing the T7 RNA polymerase were transfected with the HA-nsp3-4-V5 encoding plasmid, NeonGreen positive cells were identified by fluorescence microscopy and selected for transmission electron microscopy. Yellow boxed regions are highlighted on the right panel and the two panels below, respectively. **(E)** Comparison of HA-nsp3-4-V5 induced DMVs (as in (D)) with DMVs induced in SARS-CoV-2 infected Calu-3 cells as determined by transmission electron microscopy. Average diameters of DMVs are given on the bottom. Diameter calculations are based on the analysis of at least 10 infected cells (as reported earlier ^25^) or ~1650 DMVs detected in transfected cells.

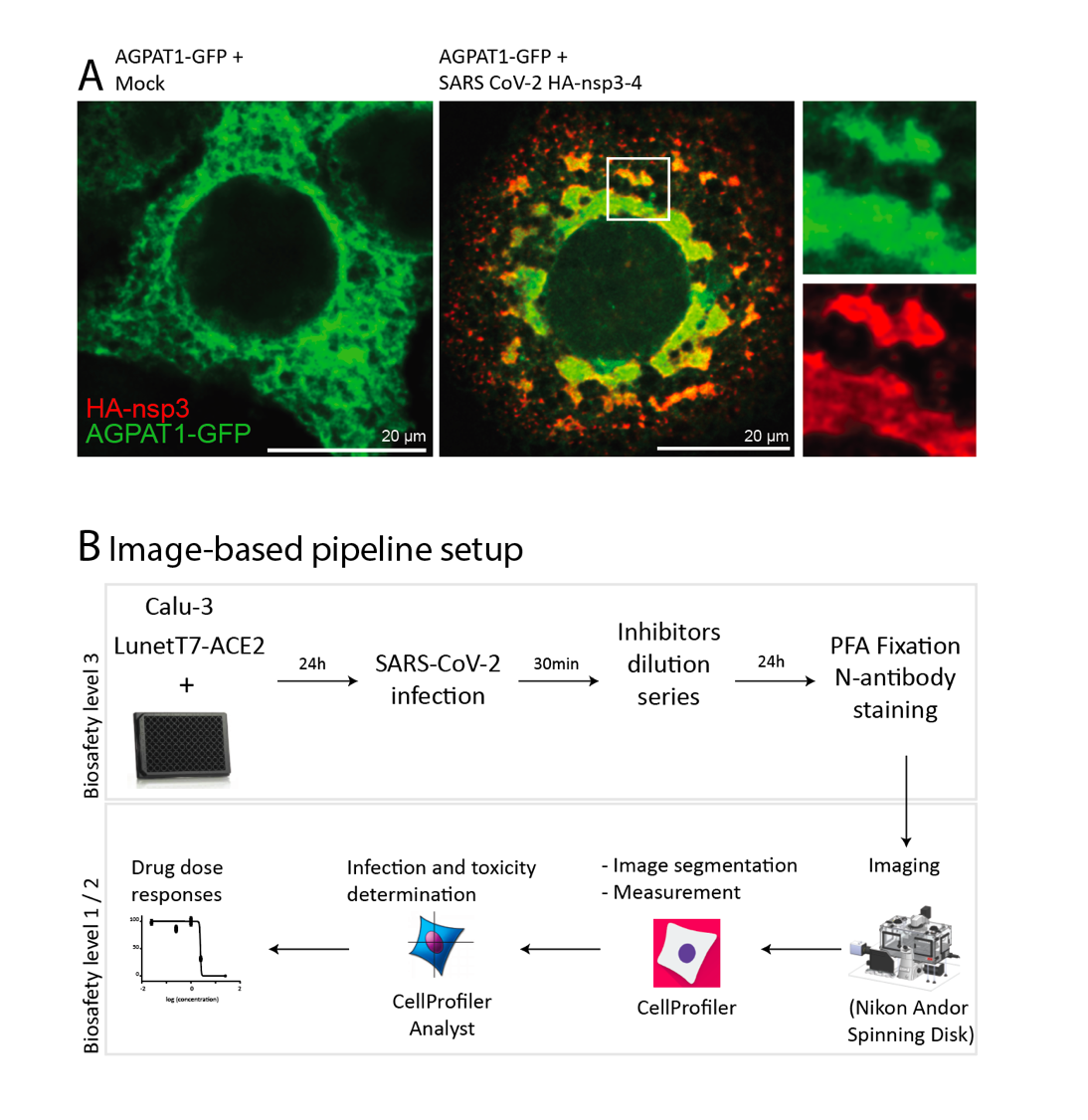

**Fig. S9. AGPATs are re-localised to SARS-CoV-2 nsp3 containing structures and imaging based pipeline to quantify viral replication. (A)** Huh7-derived cells were co-transfected with AGPAT1-GFP and SARS-CoV-2 HA-nsp3-4-V5 encoding plasmids. After 48 h cells were stained with HA-specific antibody and analyzed by confocal microscopy to visualize HA-nsp3 and AGPAT1-GFP. White square indicates the area shown as magnification on the right. (**B**) Schematics of the image-based approach used for the determination of SARS-CoV-2 replication and spread. In some experiments, infected cells were treated with drugs as indicated.

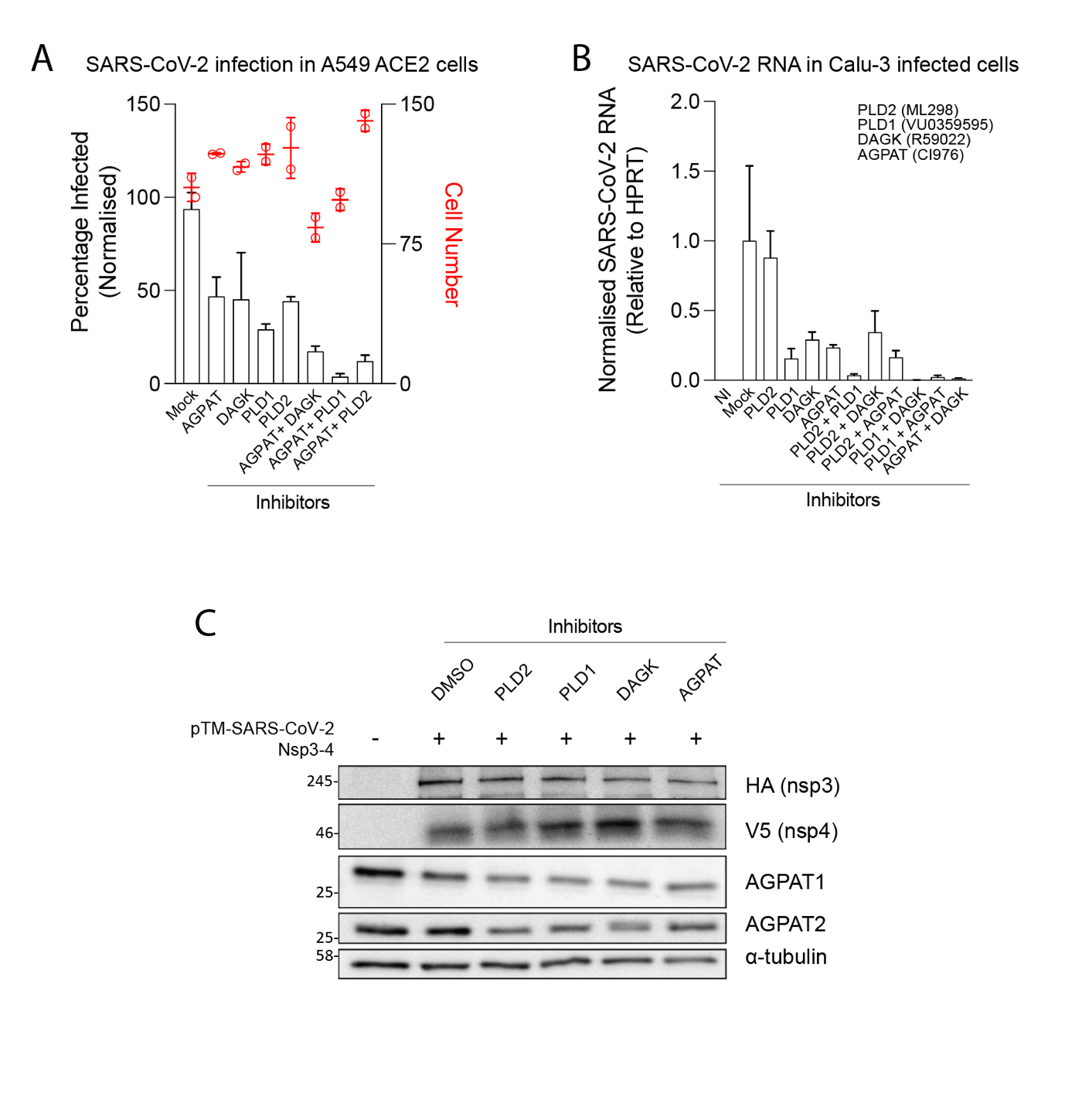

**Fig. S10. Alternative PA production pathways contribute to SARS-CoV-2 replication. (A)** A549 cells stably expressing ACE2 were infected with SARS-CoV-2 (MOI=5) and incubated with AGPAT, PLD1/2, or DAGK inhibitors, or given combinations thereof, using concentrations corresponding to IC50 values. Cells were fixed 24 h after infection and stained with a nucleocapsid-specific antibody, followed by immunofluorescence microscopy and CellProfiler based analysis. Data is normalized to the Mock, which on average has ~35% infected cells. (**B**) Calu-3 cells were infected with SARS-CoV-2 (MOI=12) for 2 h, followed by incubation with AGPAT, PLD1/2, or DAGK inhibitors using concentrations corresponding to IC50 values. Total RNA was harvested 6 h after infection and SARS-CoV-2 RNA was quantified relative to cellular HPRT mRNA using RT-qPCR. Higher MOI than **(A)** was used since (i) unbound virus was washed away after 2 h and (ii) viral replication was measured at 6 h rather than 24 h after infection. (**C**) PA pathway inhibitors do not affect SARS-CoV-2 nsp3-4 expression or processing. Huh7-Lunet/T7 cells were transfected with plasmid pTM-nsp3-4-2A-mNG encoding SARS-CoV-2 HA-nsp3-4-V5 and fluorescent NeonGreen. After 4 h, inhibitors specified on the top were added. Twenty-four hours later, total cell lysates were collected and expression levels of indicated proteins were analyzed by western blotting. α-tubulin served as loading control. These data belong to the results shown in Fig. 4E.

Table S1. Plasmids used in this study

| Plasmid Name | Backbone | Epitope tag/  reporter | Selection (E.coli/mammalian cell) | Reference |
| --- | --- | --- | --- | --- |
| pFK-JcR2a-δg (JcR2a) | pFK, derived from pBR322 | *Renilla* Luciferase | Amp | *^26^* |
| pFK-Jc1-δg (Jc1), | pFK, derived from pBR322 |  | Amp | *^27^* |
| pFK_i389LucNS3-3′_JFH1_δg (genotype 2a) | pFK, derived from pBR322 | *Firefly* Luciferase | Amp | *^28^* |
| pFK_i389Luc_NS3-3′JFH1δg (genotype 2a) (NS5A-mcherry) | pFK, derived from pBR322 | NS5A-mCherry/*Firefly* Luciferase | Amp | *^21^* |
| pTM NS3-5B | pTM1-2 |  | Amp | *^29^* |
| pTM NS3-5B (NS5A-mcherry) | pTM1-2 | NS5A-mCherry | Amp | *^21^* |
| pWPI-EGFP-CT | pWPI | EGFP | Amp/Blasti | This study. |
| pWPI-EGFP-NT | pWPI | EGFP | Amp/Blasti | This study. |
| pWPI-mCherry-NT | pWPI | mCherry | Amp/Blasti | This study. |
| pWPI-AGPAT1-EGFP | pWPI-EGFP | AGPAT1-EGFP | Amp/Blasti | This study. |
| pWPI-AGPAT2-EGFP | pWPI-EGFP | AGPAT2-EGFP | Amp/Blasti | This study. |
| pWPI-EGFP-2xPABD | pWPI-EGFP | EGFP-2xPABD (Spo20p) | Amp/Blasti | This study. |
| pWPI-EGFP-2xPABD_4E | pWPI-EGFP | EGFP-2xPABD_4E (Spo20p) | Amp/Blasti | This study. |
| pGEX-PABD | pGEX-6P-1 | GST-1xPABD (Spo20p) | Amp | This study. |
| pGEX-PABD_4E | pGEX-6P-1 | GST-1xPABD_4E (Spo20p) | Amp | This study. |
| pWPI-mCherry-Parkin | pWPI-mCherry | mCherry-Parkin | Amp/Blasti | This study. |
| pWPI-T7-Zeo | pWPI |  | Amp/Zeo | ^30^ |
| pCMV-dR8.91 | pCMV |  | Amp | kind gift from Dr. Didier Trono |
| pMD2.G |  |  | Amp | kind gift from Dr. Didier Trono |
| lentiCRISPR v2 | lentiCRISPR v2 |  | Amp/Puro | Addgene |
| lentiCRISPR_blasticidin | lentiCRISPR v2 |  | Amp/Blasti | *^3^* |
| lentiCRISPR_AGPAT1 #7 | lentiCRISPR v2 |  | Amp/Puro | This study. |
| lentiCRISPR_AGPAT1 #8 | lentiCRISPR v2 |  | Amp/Puro | This study. |
| lentiCRISPR_AGPAT2 #1 | lentiCRISPR_blasticidin |  | Amp/Blasti | This study. |
| lentiCRISPR_AGPAT2 #2 | lentiCRISPR_blasticidin |  | Amp/Blasti | This study. |
| pWPI-AGPAT1_sgRNA-resistant_wild-type (WT) | pWPI |  | Amp/Blasti | This study. |
| pWPI-AGPAT1_sgRNA-resistant_H104A, D109N (M1) | pWPI |  | Amp/Blasti | This study. |
| pWPI-AGPAT1_sgRNA-resistant_E178Q, R181A (M2) | pWPI |  | Amp/Blasti | This study. |
| pWPI-AGPAT2_sgRNA-resistant_wild-type (WT) | pWPI |  | Amp/Puro | This study. |
| pWPI-AGPAT2_sgRNA-resistant_H98A, D103N (M1) | pWPI |  | Amp/Puro | This study. |
| pWPI-AGPAT2_sgRNA-resistant_E172Q, R175A (M2) | pWPI |  | Amp/Puro | This study. |
| pTM-EGFP-PABD-Raf1_WT | pTM1-2 | EGFP-PABD_WT (Raf1) | Amp | This study |
| pTM-EGFP-PABD-Raf1_4E | pTM1-2 | EGFP-PABD_4E (Raf1) | Amp | This study |
| pWPI-EGFP-PABD-Raf1_WT | pWPI | EGFP-PABD_WT (Raf1) | Amp/Blasti | This study |
| pWPI-EGFP-PABD-Raf1_4E | pWPI | EGFP-PABD_4E (Raf1) | Amp/Blasti | This study |
| pcDNA3.1-SARS-CoV-2_HA-Nsp3-4-V5 | pcDNA3.1 | HA-Nsp3  Nsp4-V5 | Amp | This study |
| pTM SARS-CoV-2_HA-Nsp3-4-V5 (IRES-mNG) | pTM1-2 | mNeonGreen  HA-Nsp3  Nsp4-V5 | Amp | This study |

Amp, amplicillin; Blasti, blasticidin; Zeo, zeocine; Puro, puromycin.

Table S2. Reagents and resources used in this study

| Reagents or Resources | Source | Identifier |
| --- | --- | --- |
| DAPI | MoBiTec | MFPCCFA-211 |
| Lipidtox | Thermo Fisher Scientific | H34477 |
| BODIPY493/503 | Thermo Fisher Scientific | D3922 |
| Oleic acid-BSA | Sigma Aldrich | O3008 |
| Fluoromount-G | Southern Biotech | 0100-01 |
| Phorbol 12-myristate 13-acetate (PMA) | Sigma Aldrich | P8139 |
| Bafilomycin A1 (BafA1) | Sigma Aldrich | B1793 |
| Valinomycin (Val) | Sigma Aldrich | V0627 |
| TransIT-LT1 transfection reagent | Mirus Bio | MIR 2305 |
| In-Fusion HD Cloning Plus | TAKARA Bio | 638909 |
| NEB Hi-Fi Assembly Kit | New England Biolabs Inc | E2621S |
| Platinum™ SuperFi™ PCR Master Mix | Thermo Fisher Scientific | 12358010 |
| IPTG | Thermo Fisher Scientific | 15529019 |
| Slide-A-Lyzer Dialysis Cassettes, 7K MWCO | Thermo Fisher Scientific | 66370 |
| streptolysin O (SLO) | Sigma Aldrich | SAE0089 |
| Creatine Kinase (CK) | Sigma Aldrich | CK-RO, 10127566001 |
| Creatine phosphate | Sigma Aldrich | CRPHO-RO, 10621714001 |
| Guanosine 5′-triphosphate sodium salt hydrate | Sigma Aldrich | G8877 |
| Pierce Anti-HA Magnetic Beads | Thermo Fisher Scientific | 88836 |
| Pierce GST Spin Purification Kit | Thermo Fisher Scientific | 16107 |
| CellTiter-Glo Luminescent Cell Viability Assay | Promega | G7570 |
| ECL plus reagent | Perkin Elmer Inc. | NEL104001EA |
| DTT | Sigma Aldrich | D0632 |
| Glycyl Glycin | Sigma Aldrich | G3915 |
| ATP | Sigma Aldrich | A2383 |
| D-Luciferin | PJK GmbH | 102111 |
| Coelenterazine | PJK GmbH | 102161 |
| Paraformaldehyde | Sigma Aldrich | 158127 |
| 25% Glutaraldehyde | Electron Microscopy Science | 16220 |
| 16% Paraformaldehyde Aqueous Solution | Electron Microscopy Science | 15710 |
| 4% Osmium Tetroxide | Electron Microscopy Science | 19150 |
| ML298 | Sigma Aldrich | SML1077 |
| VU0359595 | Sigma Aldrich | SML0566 |
| FIPI Hydrochloride Hydrate | Sigma Aldrich | F5807 |
| R59022 | Sigma Aldrich | D5919 |

Table S3. Antibodies used in this study

| Primary antibody | Source | Identifier |
| --- | --- | --- |
| rabbit anti-HCV NS4B polyclonal antibody | *^7^* | - |
| mouse anti-HCV NS5A monoclonal antibody | *^7^* | - |
| rabbit anti-AGPAT1 polyclonal antibody | Atlas Antibodies | HPA073355 |
| rabbit anti-AGPAT2 monoclonal antibody | Cell Signaling | 14937 |
| mouse anti-alpha-tubulin monoclonal antibody | Sigma Aldrich | T5168 |
| mouse anti-beta-actin monoclonal antibody | Sigma Aldrich | A5441 |
| rabbit anti-LC3 polyclonal antibody | MBL | PM036 |
| mouse anti-HA monoclonal antibody | Sigma Aldrich | H3663 |
| rabbit anti-HA polyclonal antibody | Thermo Fisher Scientific | PA1-985 |
| mouse anti-V5 monoclonal antibody | GeneTex | GTX628529 |
| rabbit anti-V5 polyclonal antibody | GeneTex | GTX117997 |
| mouse anti-GST monoclonal antibody | Santa Cruz | sc-138 |
| mouse anti-Nucleocapsid antibody | Sino Biological | 40143-V08B |
| mouse anti-GFP antibody | Clontech | 632381 |
| mouse anti-Flag antibody | Sigma Aldrich | F1804 |

| Secondary antibody | Source | Identifier |
| --- | --- | --- |
| Goat anti–rabbit IgG-HRP | Sigma Aldrich | A6154 |
| Goat anti–mouse IgG-HRP | Sigma Aldrich | A4416 |
| Alexa Fluor 488 donkey anti-rabbit IgG | Thermofisher | A-21206 |
| Alexa Fluor 488 donkey anti-mouse IgG | Thermofisher | A-21202 |
| Alexa Fluor 488 donkey anti-mouse IgG2a | Thermofisher | A-21131 |
| Alexa Fluor 568 donkey anti-rabbit IgG | Thermofisher | A-10042 |
| Alexa Fluor 568 donkey anti-mouse IgG | Thermofisher | A-10037 |
| Alexa Fluor 568 donkey anti-mouse IgG1 | Thermofisher | A-21124 |

Table S4. Cell lines used in this study

| Descriptor | Parental cell line | resistance | reference |
| --- | --- | --- | --- |
| HEK-293T | HEK-293T |  | *^31^* |
| Huh7.5 | Huh7.5 |  | *^32^* |
| Huh7-Lunet | Huh7-Lunet |  | *^33^* |
| Huh7.5/Fluc | Huh7.5 | Geneticin | *^26^* |
| Huh7-Lunet/T7 | Huh7-Lunet | Zeocin | *^29^* |
| Huh7-Lunet/CD81H | Huh7-Lunet | Geneticin | *^34^* |
| Huh7-Lunet/subgenomic replicon [HCV wt] | Huh7-Lunet | Geneticin | *^7^* |
| Huh7-Lunet/subgenomic replicon (sg4B HA31R) [HCV 4BHA] | Huh7-Lunet | Geneticin | *^7^* |
| Huh7-Lunet/calnexin^HA^ [CNX^HA^] | Huh7-Lunet | Geneticin | *^7^* |
| Huh7-Lunet/AGPAT1-EGFP | Huh7-Lunet | Blasticidin | This study. |
| Huh7-Lunet/AGPAT2-EGFP | Huh7-Lunet | Blasticidin | This study. |
| Huh7.5/AGPAT1_sgRNA-resistant_wild-type (WT) | Huh7.5 | Blasticidin | This study. |
| Huh7.5/AGPAT1_sgRNA-resistant_H104A, D109N (M1) | Huh7.5 | Blasticidin | This study. |
| Huh7.5/AGPAT1_sgRNA-resistant_E178Q, R181A (M2) | Huh7.5 | Blasticidin | This study. |
| Huh7.5/AGPAT2_sgRNA-resistant_wild-type (WT) | Huh7.5 | Puromycin | This study. |
| Huh7.5/AGPAT2_sgRNA-resistant_H98A, D103N (M1) | Huh7.5 | Puromycin | This study. |
| Huh7.5/AGPAT2_sgRNA-resistant_E172Q, R175A (M2) | Huh7.5 | Puromycin | This study. |
| Huh7-Lunet/T7/mCherry-Parkin | Huh7-Lunet | Zeocin, Blasticidin | This study. |
| Huh7-Lunet/CD81H/mCherry-Parkin | Huh7-Lunet | Geneticin | This study. |
| Huh7-Lunet/T7/Control KO | Huh7-Lunet | Zeocin, Puromycin | This study. |
| Huh7-Lunet/T7/AGPAT DKO | Huh7-Lunet | Zeocin, Blasticidin, Puromycin | This study. |
| Huh7-Lunet/T7/ACE2 | Huh7-Lunet | Zeocin, Blasticidin, Puromycin  Neomycin | This study. |
| A549-ACE2 | A549 | Neomycin | ^25^ |
| Calu-3 | Calu-3 |  | ^25^ |

Table S5. Vectors and guide RNAs used to generate AGPAT KO cells

| Expression plamid | Target gene | Guide RNA |
| --- | --- | --- |
| lentiCRISPR_AGPAT1 #7 | AGPAT1 | 5-GGGGCTGCAGAACCACAGGG-3 |
| lentiCRISPR_AGPAT1 #8 | AGPAT1 | 5'-GAGGGAACGAGAAACCACAA-3' |
| lentiCRISPR_AGPAT2 #1 | AGPAT2 | 5'-CGCGGCCGAGTTCTACGCCA-3' |
| lentiCRISPR_AGPAT2 #2 | AGPAT2 | 5'-CTTTTACGGGCTCCGCTTCG-3' |

Data S1. Proteome analysis

LC-MS/MS RAW files obtained for untagged WT-, NS4BA- and CNX-associated membrane samples were analyzed with Proteome Discoverer 2.2 (PD2.2) using 1% false discoverer rate (FDR) on peptide-spectrum-match (PSM) and protein level. Label-free quantitation with ‘match between runs’ function was used for calculating abundance values (see *PD2.2 MS data analysis*). Results were used for ‘Significance Analysis of INTeractome’ (SAINT) using untagged NS4B as technical negative control (see *NS4B_saint_result* and *CNX_saint_result*). All proteins considered as interactors by SAINT (AvgP>0.95) of at least one of the baits (NS4B or CNX) were normalized to equal total abundance in each sample and analyzed for differential abundance between the NS4B and CNX pulldowns using the limma R software package (see *limma_output*). NS4B significantly enriched proteins (see *limma_NS4BA_enriched*) were used to generate functional networks utilizing the ClueGO v2.5.5 app embedded in Cytoscape 3.7.2 (see *ClueGO-NS4BA*).

Data S2. SiRNA screening

The following information is listed in different excel sheets: the “control siRNAs” sheet provides siRNA sequences and product numbers of siRNAs that were used in the screen; the “siRNA list_Silencer select” sheet provides all target genes and siRNAs used in the screen; the “Plate Layout” sheet shows the layout design for each plate that was used in the screen (three different siRNAs per target gene; control siRNAs were randomly placed in different plates to avoid position effects); the “A) Result_raw counts” sheet lists the luciferase counts after subtraction of the background values; in the “B) Result_sorted by z-score” sheet data are sorted by z-score; the “C) Result_gene_z-score” sheet lists the genes according to z-score after statistical analysis using data from three different siRNAs and 5 repetitions.

Data S3. Lipidome analysis

Raw data set of lipidome analysis of HCV-induced DMVs. Quantified lipids are listed including concentrations (in µM) of all individual lipid species.
